# Pollinator-relevant floral traits impact bidirectional hybridisation in the orchid genus *Gymnadenia*

**DOI:** 10.1101/2025.03.20.644302

**Authors:** Mikhaela Neequaye, Roman T. Kellenberger, Rebecca Collier, Katherine E. Wenzell, Pirita Paajanen, Rea Antoniou-Kourounioti, Lionel Hill, Hannah Gunn, Philipp M. Schlüter, Kelsey J.R.P. Byers

## Abstract

Reproductive isolation is mediated by ecological, phenotypic, and genetic barriers. Where such barriers are not upheld, hybridisation can occur, revealing insight about the formation of species. Here, we provide an integrative investigation of hybrid formation between the orchids *Gymnadenia conopsea* and *G. rhellicani* and potential isolating barriers in zones of overlap in the European Alps. We examine opportunities for hybridisation by quantifying hybrid frequency and pollinator assemblages of parent species. Next, we characterise floral phenotypes and genetic parentage of hybrids to examine how genetic inheritance of traits shapes ecological interactions, and thus fitness, of hybrids. We find that pollinator assemblages of parents overlap partially, consistent with observed rare hybrids (<1% of the population). Phenotypically, hybrids were intermediate across floral traits, consistent with all hybrids being identified as F1s, resulting from a primary hybridisation event between *G. conopsea* and *G. rhellicani*. No evidence of backcrossing or introgression was found, but direction of pollen transfer varied across sites. Additionally, hybrids were heterozygous for parental alleles at two loci impacting anthocyanin production, suggesting hybrids’ intermediate floral colour reflects inheritance of these alleles. Finally, we find little support that ploidy variation or hybrid seed infertility explains the absence of advanced generations of hybrids. Thus, we suggest that floral isolation represents an important isolating mechanism between *G. conopsea* and *G. rhellicani* in limiting the formation of uncommon, unfit hybrids and shaping species boundaries.

## Introduction

Reproductive isolation is a critical process in evolutionary biology, as it shapes (and often defines) the formation of species. Reproductive isolation may occur at several stages in reproduction, from before mating (referred to as pre-mating barriers), before the formation of a zygote (post-mating, prezygotic barriers), or after the formation of a zygote (postzygotic barriers; Coyne & Orr, 2004). The processes leading to the formation of these barriers, and the order in which various barriers may arise, remains a fundamental question in evolution. Hybridisation occurs when species overcome barriers to reproduction, thus highlighting a key opportunity to study the formation and strength of reproductive isolation. Widespread evidence of hybridisation in wild populations continues to grow (Monnet et al., 2025), alongside the application of novel genetic tools for characterising individual species and hybridisation events (Goulet et al., 2017). Where hybridisation occurs infrequently in populations, a natural laboratory forms in which isolation barriers in flowering plants can be studied.

For many flowering plants, reproduction depends on the movement of pollen via animal pollinators, and thus pollinators may represent a vital source of pre-mating reproductive isolation, which can be among the strongest barriers – especially in sympatry or overlapping ranges (Lowry et al. 2008; Ollerton et al., 2011; Ramsey et al., 2003). Therefore, studying pollination helps us understand species formation and how reproductive isolation is maintained in sympatry and secondary contact (Coyne & Orr, 2004). Pollinator isolation may be achieved through the establishment of suites of floral traits often associated with specific pollinator guilds (Fenster et al., 2004). For closely related species, adaptation of floral traits to distinct pollinator guilds can provide an essential means for maintaining pre-zygotic isolation in areas of sympatry (Bradshaw & Schemske, 2003; Jones, 1978; Katzer et al., 2019; Wessinger, 2021). In cases where hybrids between parapatric species in secondary contact result in fitness costs, prezygotic isolation resulting from pollinator behavioural responses to divergent floral traits has been shown to result in reinforcement (Hopkins and Rausher 2012). This illustrates the important links between floral trait variation among parapatric species, pollinator responses, and the formation or reinforcement of reproductive isolation. The ultimate evolutionary outcome of establishing these dynamics is pollinator-mediated assortative mating, which is a key pre-zygotic isolating barrier in flowering plants (Christie et al., 2022; Waser, 1993). However, these pre-zygotic barriers are not always upheld, which can lead to “leaky” reproductive isolation and hybridisation between distinct species (Jersáková et al., 2010).

Due to their high species diversity and specialised floral phenotypes and pollinator interactions, orchids (Orchidaceae) offer compelling opportunities to investigate reproductive isolation, making them models for pollinator-mediated plant diversification (Schiestl & Schlüter 2009). The *Gymnadenia* orchids of the European Alps provide a valuable model in which to study the evolutionary ecology of hybridisation due to evidence of pollinator-mediated selection on floral traits and emerging knowledge connecting these traits to their genetic basis (Kellenberger et al., 2019; Gupta et al., 2014). The *Gymnadenia* orchids display a wide range of floral morphologies and attractive traits, are closely related (Piñeiro Fernández et al., 2019) and produce hybrids (Hedrén et al., 2018). *Gymnadenia conopsea* and *G. rhellicani* hybridise to form the purported nothospecies (i.e., a hybrid species formed through crossing of two distinct parent species) *G. x suaveolens* in their limited zones of sympatry. However, hybrids of *G. conopsea* and *G. rhellicani* are thought to be rare (Hedrén et al., 2018) and questions remain regarding the genetic evidence for their parental origins and degree of hybridisation. Hybrid scarcity is suggested to be a result of the distinctive floral morphology, and therefore pollination systems, of the closely related *Gymnadenia* parent species (Pridgeon et al., 1997; Vöth, 2000). Hybrids therefore offer a natural laboratory in which to examine reproductive isolation - or lack thereof - among parent species. Studying hybrids can further our understanding of the biochemical and genetic mechanisms underlying pollinator-relevant floral traits (defined as those shaping pollinator fit and/or preference) likely to impact reproduction and fitness.

*Gymnadenia x suaveolens* hybrids and their proposed progenitors possess distinctive floral traits, many of which have been shown to be under selection from insect pollinators in at least one parent (Sletvold & Ågren, 2010, Chapurlat et al., 2015; Kellenberger et al. 2019; Chapurlat et al., 2020). Despite some overlap in observed pollinator families, particularly among Lepidoptera, effective pollinator composition differs between the two parent species *G. conopsea* and *G. rhellicani* (Claessens & Kleynen, 2011; Delforge, 1994, 2006; Hackney, 2007; Sun et al., 2014; Sun et al., 2015; Vöth, 2000). This is proposed to be due in part to their variation in nectar spur length, which coincides with the average proboscis length of effective pollinators for *G. conopsea*, and is expected for *G. rhellicani* as well (Schiestl & Schlüter, 2009; Sletvold & Ågren, 2010). Variation in nectar spur length aligns with longer spurs conferring higher nectar volumes (Brzosko & Mirski, 2021), reflecting increased floral rewards for larger pollinators. Hedrén et al., 2018 propose hybridisation to occur most frequently in one direction (*G. rhellicani* paternal to *G. conopsea* maternal) based on pollinia placement on the probosces of pollinators due to these mechanical isolation mechanisms. However, the genetic identity of *G. x suaveolens* has yet to be characterised.

Regarding pollinator responses to other floral traits, earlier work has examined physiological responses in pollinators to floral scent in *G. conopsea* (which at the time included *G. densiflora,* now elevated to species rank) and the closely related species *G. odoratissima* (Huber et al., 2005). Moreover, *G. conopsea* has become an important example of pollinator-mediated selection on floral scent (Chapurlat et al., 2019; Chapurlat et al., 2018; D’Auria et al., 2020) and morphology (Sletvold & Ågren, 2010; Chapurlat et al., 2015). Despite evidence that pollinators influence floral traits in this species, a recent study concluded that pollinator sharing in *G. conopsea* and *G. densiflora* does not lead to diversification of scent via character displacement (Joffard et al., 2022), as would be expected under pollinator-mediated premating isolation. Nonetheless, pollinator sharing is unsurprising in this system given the similarity in floral morphology and colour of these two species, as evidenced by their status as a single species until recently (Marhold et al., 2005; Stark et al., 2011; Chapurlat et al., 2020). Thus, expectations of how pollinators may shape the likelihood of interspecific mating (i.e., hybridisation) will differ for species pairs that vary in floral signals, such as *G. conopsea* and *G. rhellicani*. Detailed characterisation of floral scent has not been performed for the *G. x suaveolens* hybrids nor for *G. conopsea* where it co-occurs with *G. rhellicani*. Furthermore, whilst attempts have been made to reconstruct the phylogenetic relationships within this group (Bateman, 2021; Brandrud et al., 2019; Piñeiro Fernández et al., 2019), thorough phenotypic characterisation alongside whole genome sequencing has yet to be performed for these species and their proposed hybrids. Consequently, our understanding of the relationship between pollinator-relevant floral traits, reproductive isolation, and hybridisation in *Gymnadenia* remains incomplete.

In the field, the *G. x suaveolens* hybrid individuals are immediately distinguishable by their moderately dense, intense magenta inflorescences, whilst one proposed progenitor parent *G. conopsea* has open lilac-coloured inflorescences, and the other parent, *G. rhellicani*, displays contrasting densely-clustered inflorescences that are typically dark red in colour (Figure 1A). These floral colours vary intra-specifically, with *G. conopsea* displaying a range from light pink to deeper purple petals (Gross & Schiestl, 2015) and the red intensity of *G. rhellicani* varying from black-red to almost white in some populations (Kellenberger et al., 2019). In *G. rhellicani*, genetic regulation of pigment production is influenced by pollinator interactions, where pollinator-mediated overdominance results in heterozygous ‘red’ floral morphs having greater fitness than homozygous ‘black’ (dark red) and ‘white’ morphs (Kellenberger et al., 2019). Kellenberger et al., (2019) demonstrate the overdominance of a non-synonymous SNP with three allelic states at position 663 in the R2R3 MYB transcription factor, MYB1, in *G. rhellicani*. The most common state is a C, with other alleles being either a G or an A, in the last exon of the coding region. This results in a premature stop codon, truncated protein and downstream reduction in the expression of target-gene ANTHOCYANIDIN SYNTHASE (ANS) in the less common alleles. This ultimately leads to a reduction in the production of cyanidins in *G. rhellicani*, resulting in a paler flower colour.

**Figure 1:**
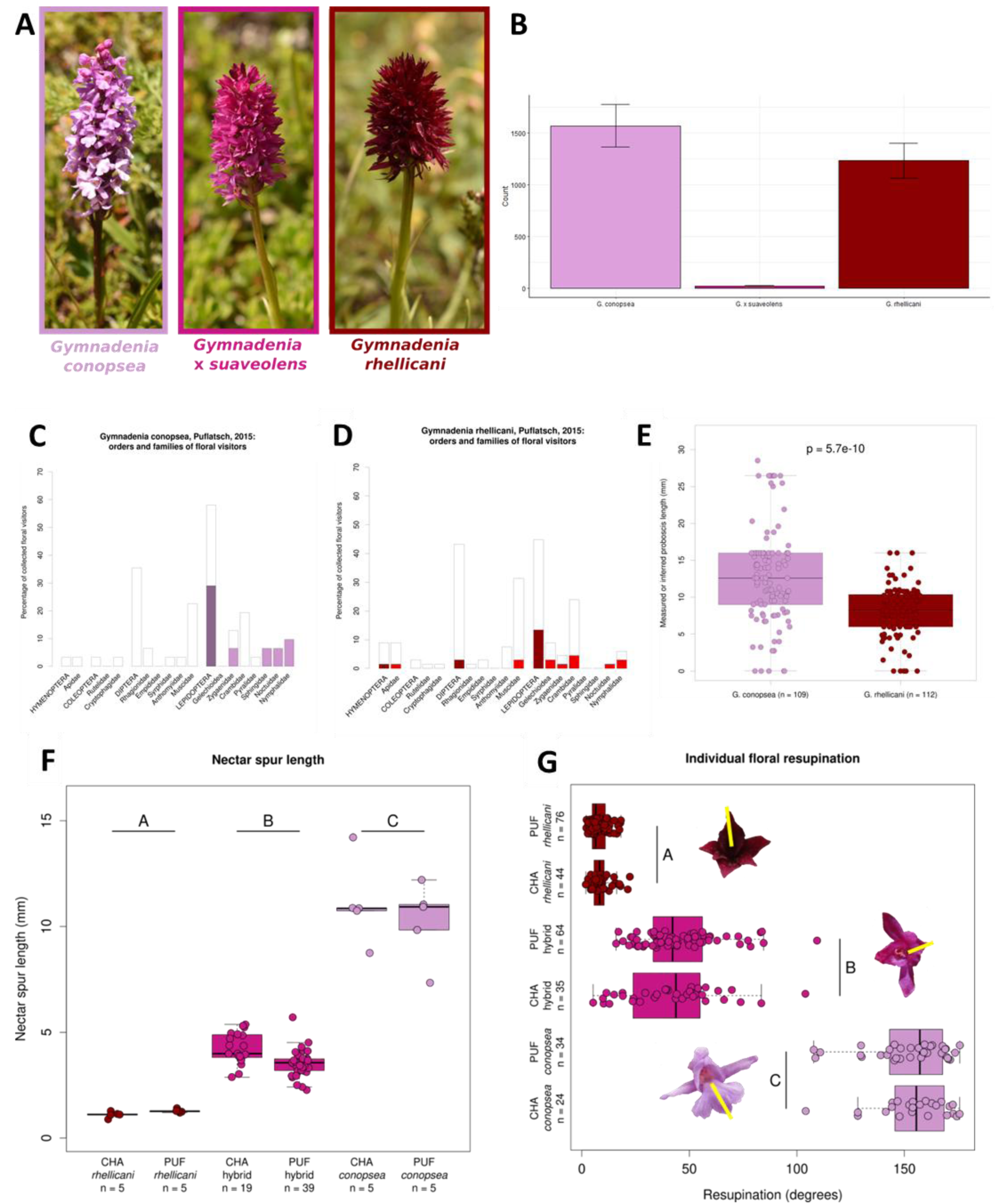
Hybridisation, pollination and pollinator-relevant morphology. (A) Focal species studied here: *G. conopsea*, *G. rhellicani*, and their putative hybrid *G. x suaveolens.* (B) Relative frequency of parents and hybrids as measured in 2015 and 2025. Floral visitors captured on *G. conopsea* (C) and *G. rhellicani* (D) in 2015 at PUF. Open bars: all visitors. Coloured sub-bars: visitors carrying orchid pollinia. Darker bars represent order total; paler bars represent family totals. (E) Differences in floral visitor proboscis lengths from the historical literature (excluding the present study). (F) Nectar spur length in millimetres for each species at each site. (G) Individual floral resupination (rotation) in degrees for each species at each site. Inset images show the average degree of rotation for each species using the yellow line marked across the labellum (largest tepal).

Thus, building on this previous work, this system allows us to investigate the connection between ecological interactions that shape mating events (i.e., pollinators), phenotypic floral traits likely to impact fitness of parents and hybrids, and their underlying genetic components, in order to shed light on the formation of reproductive barriers in a megadiverse family. In this study, we investigate the extent of hybridisation between co-occurring populations of *Gymnadenia conopsea* and *G. rhellicani* under secondary contact. First, we quantify the relative frequency of hybrids to shed light on the apparent strength of isolating barriers between parent species. Because the formation of hybrids relies on interspecific pollen transfer mediated by pollinators, we then explore the pollinator assemblages of both parent species to understand the degree of overlap and opportunities for shared pollinators to contribute to interspecific pollen transfer and therefore hybridisation. We then characterise the hybrids in depth by quantitatively phenotyping putative hybrids and parents across a suite of floral traits known to influence pollinator interactions, to investigate the nature of hybrid phenotypes and likelihood of floral isolation. Further, we use whole genome sequencing to investigate the genomic evidence of parentage and whether putative hybrids based on phenotype in fact represent F1 hybrids. Additionally, we use these data to assess evidence of subsequent backcrossing or advanced generation of hybrids, informing the strength of current isolating mechanisms in natural populations. Finally, we explore the relationship between hybrid parentage and observable phenotype in floral traits by examining genotypes at two loci influencing anthocyanin production, which are known to impact floral colour in the parent species. We then consider possible barriers to reproduction between parent species that may act to limit hybridisation in this system.

## Materials & Methods

### Study species & Field Sites

The study species *Gymnadenia conopsea* (L.) R.Br., *G. rhellicani* (Teppner & E. Klein) Teppner & E. Klein and their proposed hybrid, the nothospecies *G. x suaveolens* (Vill.) Rchb. f. are fragrant, pollinator-rewarding orchids, found in European meadows and grasslands from Scandinavia (Olofsson, 2021) to the Alps and Mediterranean region (Claessens & Kleynen, 2011; Kellenberger et al., 2019). These orchids can be observed flowering from mid-May until mid-August (Delforge, 1994, 2006; Gross & Schiestl, 2015; Hackney, 2007) with flowering times becoming increasingly variable due to abiotic factors like temperature fluctuations, as a result of rapidly changing climate (Kolanowska et al., 2021). Both *G. conopsea* and *G. rhellicani* are generally diploids with a chromosome number of 2n=40 (Rice et al., 2015), though polyploidy has been reported in *G. conopsea* (Stark et al., 2011). These fragrant orchids are perennials that produce a single inflorescence per plant per flowering year but do not necessarily reflower every year. These inflorescences are cylindrical in *G. conopsea* and conical to spherical in *G. rhellicani*, with variability between these two extremes observed in *G. x suaveolens*.

The two proposed parent species of *G. x suaveolens* are only sympatric in the lower elevational edge of the range of the more alpine *G. rhellicani* in the European Alps. For this reason, we chose two field sites where both species grow in sympatry and *G. x suaveolens* has been reliably observed and documented (Supplementary Figure 1; Claessens & Kleynen, 2016; Delforge, 1994, 2006): Puflatsch (PUF), Seiser Alm, Südtirol, Italy (2125m) and Chandolin (CHA), Kt. Wallis, Switzerland (2225m), with the two sites separated by ca. 300km. Sampling was conducted in 2014-2016 and 2021-2023 for floral trait and fitness data. In 2015, we counted individuals of each species along a transect at PUF to quantify the relative abundance of the *Gymnadenia* species and hybrids (Kellenberger et al., 2019; site “RTK3”) and repeated in 2025 to assess change over time. The transect was started at the highest point of the site and a compass was used to walk a straight line across the site, counting each orchid within one metre either side of the transect. When the edge of the site was reached, the transect was continued from approximately two metres downslope. This method was continued until the whole site had been covered for a whole population count of the study organisms, and the percentage frequency of hybrids was compared across sampling years. Transect data were plotted in R (R Core Team, 2025) using the package ggplot 2 (Wickham, 2016).

### Potential pre-mating barriers

#### Pollinator ecology

In 2015, floral visitors at site PUF were collected across a period of 2 days including morning, afternoon, and evening (dusk and full night) collections. Floral visitors were noted to functional group and time of day (diurnal or nocturnal) when observed landing and interacting with an orchid with open flowers. Where possible, visitors were captured using a net and moved into 50 ml falcon tubes with ethyl acetate killing fluid for later identification. Visitors were assessed for the presence of pollinia, to distinguish which visitors are likely to be capable of effectively pollinating orchid flowers. To supplement these observations, a literature search was undertaken to determine identities of previously recorded visitors to the two putative parent species. Data on pollinator and visitor proboscis lengths for both 2015 collections and the historical visitation data was taken from the literature for captured species where available (see Table S1). When species-level data were unavailable, an average value for the same genus (or for family if a genus value was unavailable) was used. Proboscis lengths of Coleoptera and non-Empididae Diptera was inferred to be 0.01mm as the proboscis of these insects is too large to fit into the quite narrow nectar spurs of either orchid species.

### Quantifying floral traits of parent species and hybrids

#### Floral morphology

##### Morphological measurements

At both field sites, total plant height and inflorescence height were measured using a Stanley 035455 2m Wooden Folding Rule. For individual flowers, display height, display width and nectar spur length were measured using 150mm digital callipers (Linear Tools, RS Components). These traits were chosen because overall size of floral display is expected to be important for pollinator attraction, and length of nectar spur directly relates to reward access and fit of pollinators, based on their respective proboscis lengths (Schiestl and Schluter, 2009).

##### Resupination

Another floral trait expected to be important for pollinator fit and pollen transfer is resupination. Resupination is the process in which the positioning of the labellum (the uppermost tepal of orchid flowers) is inverted, or twisted, during development to form a landing platform for pollinators (Cardoso et al., 2024). It is a gravitropic mechanism regulated by auxin (Yam et al., 2009), where buds develop and twist, so the labellum is positioned at the bottom of the flower, typically around 180° to form a platform for floral visitors (Arditti, 2003; Zhang et al., 2018), though the degree of resupination can vary among orchid species. To measure resupination of parent species and hybrids, whole inflorescences were photographed in the field. In GIMP (Team, 2019), a line was drawn down the inflorescence on the image using the measurement tool and the image was adjusted using a rotation tool to an angle of 180° . The angle of resupination was then measured by drawing a line from the labellum to the centre of the flower. This was repeated for three flowers per inflorescence and a mean angle of resupination was then calculated for each plant.

#### Floral scent

Floral scent is known to play a key role in the attraction of pollinators of *Gymnadenia* orchids, with specific volatiles demonstrating both physiological and behavioural activity (Chapurlat et al., 2019; Huber et al., 2005). Thus, we examined floral scent using dynamic headspace capture (Raguso & Pellmyr, 1998) on single individuals in the field as follows. For 2021-2023 samples, intact inflorescences were enclosed in a PET oven bag (ca. 1L volume; Toppits, Minden, Germany) attached to a battery-driven pump (Spectrex PAS 500, Spectrex Corp., Redwood City, CA, USA) connected to a volatile trap consisting of a trimmed Pasteur pipette loaded with silanized glass wool and 100mg PorapakQ. Air was pulled from the bag through the trap at a flow rate of 100-200 ml min−1 for 40 minutes in earlier sampling seasons (2021-2022) and increased 2 hours from 2023 for logistical purposes. The traps were kept cool in the field using ice packs prior to elution from each column using 600 µl of 10% acetone in HPLC-grade hexane and then stored in the freezer at -4°C. Background VOC levels were determined with simultaneous samples from the same time and location in field using empty bags. Samples were concentrated 3-fold from a sample aliquot of 150 µl to 50µl under ambient room air and 3µl was injected using a Gerstel MPS system into an Agilent 7890B GC with 5977A/B MS GC-MS in splitless mode with the following method. A Zebron ZB-5HT w/GUARDIAN column (Phenomenex, Macclesfield, UK; 35 m, 0.25 mm, 0.1 μm) was used, and helium was used as the carrier gas at a constant flow of 1 cc min−1. The initial oven temperature was 50°C for 4 min, followed by a heating gradient of 5°C min−1 to 230°C, which was then held isothermally for 4 min. The MS was operated in scan mode (50-600m/z). Initial identification and quantification of compounds was performed with Agilent Unknowns software (v10.1, Agilent, Santa Clara, CA, USA) based on a mass spectral library built on calibration curves using three to five different concentrations of authentic reference standards. This was followed by a custom analysis pathway for removal of contaminants identified in the blank columns collected and quantification of all the remaining compounds (as described in (Wenzell & Neequaye et al., 2025)). Samples from 2021, 2022 and 2023 which were collected as above were pooled with samples from 2014-2016 which were collected, run, and analysed as in (Kellenberger et al., 2019). As samples were pooled, sampling year was factored into the data presented and all statistical analyses.

#### Floral pigment

##### Reflectance spectrophotometry

Floral colour, another trait known to impact pollinator visitation in this system (Kellenberger et al., 2019), was quantified based on both in vivo reflectance and extraction pigments. Samples collected from 2014-2016 were measured and analysed as in (Kellenberger et al., 2019). For 2021, measurements were made on a spectrophotometry set up including an Ocean Insight FLAME-S-UV-VIS-ES Assembly with a Pulsed Xenon Lamp at 220 Hz, 220-750 nm with a Premium 600 um Reflectance Probe using Ocean View spectroscopy software, Ocean Insight. The labellum of fresh flowers, taken from the field on the day, were used for reflectance spectrophotometry, with three technical replicates per flower and three flowers per plant. The entire setup was enclosed in a black velvet cloth to block ambient light. A white standard (Certified Reflectance Standard, Labsphere) and electric dark were used to calibrate the spectrophotometer readings prior to each set of measurements.

##### Pollinator perception

To characterise how floral colours may be perceived by pollinators, we plotted the reflectance data of labella on several models of pollinator visual systems. For bees, we used trichromatic models of bee (*Apis mellifera*) visual systems using R v.4.2.287 with R package pavo version 2.7.1109, which estimates level of contrast of a colour signal against a vegetative (green) background and displays the results on the bee colour perception hexagon. In addition, we used the pavo package to characterise how floral colours may be perceived by tetrachromat pollinators (including birds and day-flying butterflies) and visualised these in a three-dimensional tetrahedral space with the vertices representing the four cone receptors.

##### Anthocyanin analysis

Anthocyanin extraction was modified from (Butelli et al., 2008). Three whole flowers per individual were preserved in absolute methanol prior to extraction to facilitate transfer from the field. Flowers were ground in 80% methanol and 1% hydrochloric acid and shaken overnight at 4°C. Samples were spun down at maximum speed with an aliquot of supernatant taken for UHPLC-MS and total spectrophotometric absorption at 525nm. Samples were run on an Agilent 1290 Infinity II UHPLC equipped with PDA and 6546 quadrupole time-of-flight (Q-ToF)-MS detector Separation was on a 100×2.1mm 2.6μ Kinetex EVO column using the following gradient of acetonitrile versus 1% formic acid in water, run at 600μL.min-1 and 40°C. The diode array detector collected individual channels at 350nm (bw 4nm) and 525nm (bw 50nm), and also full spectra from 220-640nm, at 10 Hz.The Q-ToF collected positive electrospray MS using the JetStream source, together with data-dependent MS2. The MS was calibrated before use, but also had two lock-masses infused during the runs, at 121.05087 and 922.009798; components of Agilent’s normal ESI calibration mix). Spray chamber conditions were 325°C gas, 10L.min-1 drying gas, 20psi nebulizer pressure, 400°C sheath gas, 12L.min-1 sheath gas, 4000V Vcap (spray voltage) with a nozzle voltage of 1000V. The fragmentor voltage was 180V.

Floral pigments are often coupled with floral scents, as similar precursor compounds and biosynthesis pathways are involved in their production. For example, anthocyanin pigments are related biosynthetically to benzenoid and phenylpropanoid volatiles (Li et al., 2024). For these reasons, we also assessed evidence for integration among these floral traits, using Pearson Correlation Coefficients (PCC), to understand their biological basis and implications for trait evolution.

### Genetic analysis

#### Flow cytometry

Variation in ploidy often functions as a barrier to hybrid formation (Ramsey & Schemske, 1998). Both *G. conopsea* and *G. rhellicani* are generally diploid with 2n=40 (Rice et al., 2015), though variation has been reported (e.g., Stark et al., 2011). To assess whether variation in ploidy level among parent species or hybrids could constitute a reproductive barrier, we measured ploidy level using flow cytometry with a protocol modified from (Doležel et al., 2007; Gross & Schiestl, 2015; Kellenberger et al., 2019). The internal standard was fresh *Phaseolus coccineus* “Scarlet Emperor” leaves, which contain a mix of diploid (2n) and tetraploid (4n) cells while the latter undergo DNA replication and cell division. For the internal standard, young leaves were chopped in a petri dish using a razor blade, and saturated in Baryani’s Solution (Otto I substitute, 0.1M citric acid monohydrate, 0.5% v/v Triton-100). The solution was then extracted, filtered through a 30µm CellTrics filter, and placed on ice. Orchid pollinia samples (which contain a mix of haploid (n) and diploid (2n) cells) were ground in a 1.5ml Eppendorf using a small pestle with 40µl of Baryani’s Solution. 5 µl of the filtered internal standard solution were added to each pollinia sample, and the solution then filtered as above. Samples were spun for 5 minutes at 2000 rpm. The resulting pellet was resuspended in 40µl of Baryani’s solution and placed on ice for 1-3 hours. Whilst chilling the pellet, 996µl Otto II solution (0.4M Na2HPO4 • 7H2O) was combined with 4µl of DAPI in an Eppendorf tube, and the tube was wrapped in aluminium foil to keep the DAPI in the dark. Samples from 2014-2016 were analysed on a Beckman Coulter Cell Lab Quanta flow cytometer (Beckman Coulter, USA) and those from 2021-2023 were analysed using a BD FACSMelody cell sorter (BD Biosciences, Wokingham, UK). The ploidy was then determined by the ratio between the median value for the internal standard (first, i.e. diploid, peak of two internal standard peaks) and pollinia (second, i.e. diploid, peak of two pollinia peaks) sample peaks displayed on each flow cytometric histogram, with values for diploids ranging from 3.29-5.53.

#### Whole Genome Sequencing

To identify the parental origins of putative hybrids, we performed whole genome sequencing of parents and hybrids derived from both sampling sites. DNA extractions were performed on silica gel-dried leaves collected from the field using the Cytiva Nucleon PhytoPure Genomic DNA Extraction Kit Code: RPN8511 (Cardiff, Wales). Samples were sent to an external sequencing provider for 150bp paired-end Illumina sequencing using a NovaSeq 6000 following library preparation using NEBNext Ultra II FS DNA Library Prep. Parent species were sequenced with 20x coverage and hybrids with 10x. This includes 10 parent individuals (5x *G. conopsea*, 5x *G. rhellicani*) and 21 hybrids from Chandolin, and 10 parents (5x *G. conopsea*, 5x *G. rhellicani*) and 42 hybrids from Puflatsch.

Because, there are no reference genomes for *G. conopsea* or *G. rhellicani*, all the whole genome level work was done reference free using k-mer methods, using the sequenced fastq files for each sample. Counts of shared sequence content between hybrids and parents was performed with an in-house pipeline incorporating yak (https://github.com/lh3/yak) software, which includes k-mer counting and parental share counting functions. Firstly the k-mers from raw fastq reads of each potential parent were counted using the ‘yak count’ function with the following parameters: k-mer size 57 and Bloomfilter 37. Then the function ‘yak triobin’ was used with the k-mer counts of the potential parents and the hybrid as the offspring. Results were printed as percentages supporting each parent with an in-house script.

Because orchids’ plastid genomes are derived from the maternal or “egg” parent (Cafasso et al., 2005; Corriveau & Coleman, 1988), plastid genome markers can be used to determine maternal parentage (Hedrén et al., 2018). Here, we used whole genome sequencing to determine hybrid parentage using both genome balance proportions (detailed above) and copy number variation of protoplastic SNPs in the same trnL intron as in (Hedrén et al., 2018). Parent species analysed were site-specific, in that hybrid trnL intronic sequences were compared to parents of that corresponding site. The raw fastq reads from the hybrids were mapped to the plastid sequences in addition to genes known to influence anthocyanin pigment phenotypes in this system using bwa v0.7.17. (Li & Durbin, 2009). Plastid sequence identity was determined using the number of *trnL* intronic repeats published in (Hedrén et al., 2018). *MYB1* and *ANS* sequences were derived from (Kellenberger et al., 2019) and visualised in iGV (Robinson et al., 2011), where the identity of the sequences were visually assigned.

### Reproductive fitness

We asked whether the hybrids experience a penalty in fitness and whether this might provide a reproductive isolating barrier. To assess reproductive fitness, we measured the following.

#### Number of potential ovaries

The number of potential ovaries is the summation of buds, open flowers, and senesced flowers counted on each plant’s single inflorescence and is used as a metric of potential individual flowers capable of producing fruit and therefore seed. The images analysed were captured over a period of 6 years (2014-2016 and 2021-2023) intermittently, from both sampling locations. They were loaded into the GIMP (GNU Image Manipulation Program) software (Team, 2019), and manually scored for buds, open flowers, and senesced flowers. This process was repeated three times per image to produce an average to control for user error. The average number of buds, flowers and senesced flowers was then multiplied by a factor of 1.22, to encompass the parts of the inflorescence which were not captured in the photo, so that each number was representative of a whole plant in accordance with similar analysis performed in Kellenberger et al., 2019.

#### Seed counts

Open-pollinated seeds were extracted from dry fruits, ranging in number from 3 to 5 fruits per individual plant (in accordance with permitting requirements, only a small subset of fruits was collected for each plant to avoid population viability risks). The seeds were carefully transferred to marked conical-bottom 2ml freezer vials. The vials were stored in a saturated lithium chloride (LiCl) atmosphere with open caps to dehydrate. The LiCl environment was created using a Tupperware box, filled 1cm deep with lithium chloride and 3cm of sterile water. After a month of drying, the seeds were spun down gently in a centrifuge at 3000 rpm. To quantify the number of seeds, the seed vials were photographed from the side at bench height. Photographs were analysed using GIMP software (Team, 2019). The volume of seeds in each vial was calculated by measuring the height and width of the seed tube using the GIMP measurement tool (Supplementary Figure 2A). Using basic geometry for the volume of a cone (bottom of the vial) and cylinder (rest of the vial), measurements were converted to cubic millimetres and from there to seed counts using the known seed volumes for *Gymnadenia* sp. (representing *G. conopsea*) and “*Nigritella* sp.” (now *Gymnadenia* subgenus *Nigritella*, representing *G. rhellicani*) from (Arditti & Ghani, 2000). Potential ovary counts (see above) were used to scale up the number of seeds per collected fruit to the number of seeds per inflorescence.

#### Seed fertility

To assess whether seeds contained a fully developed, filled-out embryo, unstained light microscopy with a Zeiss Axio Zoom V16 (Zeiss, Cambourne, UK) was used to determine the fertility of individual seeds. After calculating their volume (see above), seeds were examined for fertility. As orchid seeds contain no endosperm (Vinogradova & Andronova, 2002), the seeds are mainly transparent under light microscopy, with a solid embryo showing as an opaque rounded shape in the centre if fertile, and no embryo present (completely translucent seed) if infertile (Supplementary Figure 2B). A petri dish was scored with grid lines 1cm apart using a black marker. The microscope was set to bright field, aperture 100%, magnification 10.5x. The Zeiss ZEN 3.1 software was paired to the microscope camera and was used to capture images of the seeds which were poured onto the scored petri dish and scattered evenly. Images were captured of each grid section, until approximately 100 seeds per sample were captured. Seeds were marked for fertility using a custom R script (‘CountSeeds_A_03.02’) (Schlüter & Byers, 2015). They were manually marked in the script as fertile or infertile seeds, using left and right mouse clicks respectively with R saving the progress after each 100 seeds marked per sample. The proportion of fertile seeds was recorded, and the number of fertile seeds per individual was calculated by multiplying the percentage of fertile seeds by the total number of seeds for that individual.

## Results

### What factors control the formation of hybrids between *G. conopsea* and *G. rhellicani*?

#### Hybrids are rare, even in large parental populations

We began our study by characterising how common hybrids are in zones of sympatry between the two parent species *Gymnadenia conopsea* and *Gymnadenia rhellicani*. In the field, *G. x suaveolens* hybrids are recognisable by their intermediate floral phenotype (Figure 1A). At PUF, we recorded 1362 *G. conopsea*, 1062 *G. rhellicani* and 19 *G. x suaveolens* hybrids in 2015, and 1773 *G. conopsea*, 1399 *G. rhellicani* and 27 *G. x suaveolens* in 2025 (Figure 1B). Although the counts for each species were higher in 2025, the proportions of each species are remarkably stable across 10 years. The hybrids made up 0.789% of the total population of orchids in 2015 and 0.844% in 2025, which suggests isolating mechanisms may limit but not entirely prevent hybridisation. Thus, we further explore the phenotypic and genetic basis of hybrids and potential barriers to understand the source and strength of reproductive isolation.

#### *G. conopsea* and *G. rhellicani* have distinct pollinator guilds with some overlap in Lepidoptera

As hybrid formation relies on interspecific pollen transfer, we assessed whether pollinators might play a role in reproductive isolation between the parent species and thus hybrid rarity. To investigate the degree of pollinator sharing between parent species, we combined historical observation data from the literature with our own field observations (Figure 1C, D, Table S1, Table S2). Our observations (Figures 1C and 1D) and the historical literature suggests Lepidopteran species as focal pollinators for both parent species, especially *G. conopsea*, where Hymenoptera and Diptera are frequently observed visitors (Claessens & Kleynen, 2011; Faegri & Van der Pijl, 1979; Proctor et al., 1996; Vöth, 1999; Vöth, 2000) (Figure 1C). Similarly, *G. rhellicani* also has Hymenopteran and Dipteran pollinators in addition to Lepidopterans (Claessens & Kleynen, 2011; Faegri & Van der Pijl, 1979; Vöth, 2000) (Figure 1D). Summarising based on broad taxonomic groups, *Gymnadenia conopsea* is mostly pollinated by Lepidoptera with long probosces and *G. rhellicani* by Lepidoptera with medium-sized to short probosces (Schiestl & Schlüter, 2009) (Figure 1E). Both *G. conopsea* and *G. rhellicani* share five families of Lepidopterans known to carry pollinia, three day-active (Pieridae, Nymphalidae and Zygaenidae) and two night-active (Noctuidae and Geometridae) (Claessens & Kleynen, 2011, 2016; Faegri & Van der Pijl, 1979; Proctor et al., 1996; Vöth, 1999; Vöth, 2000). From our field collections of co-occurring parent species at PUF, we found that the only overlapping families of validated pollinia-carrying pollinators between the two species were Noctuidae, Nymphalidae, and Zygaenidae, and only *Zygaena exulans* overlapped at the species level. This is consistent with expectations based on the long nectar spurs of conopsea, which are associated with access to nectar rewards being limited to only pollinators with long probosces. We therefore conclude that interspecific pollen transfer is likely possible between parent species due to their sharing of a subset of effective pollinators.

### How do phenotypes of hybrids compare to parent species?

#### Hybrid morphology is intermediate

When looking at overall morphological variation between the proposed parent species, *Gymnadenia conopsea* and *Gymnadenia rhellicani,* and their hybrids, we found that *G. conopsea* and *G. rhellicani* differ in total plant height, with hybrids being intermediate (Supplementary Figure 3A). This is also found in inflorescence height (Supplementary Figure 3B). Looking closer at inflorescences, flower number (calculated as the total number of potential ovaries, i.e. buds plus open flowers plus senesced flowers) is found to be the same between *G. conopsea* and *G. rhellicani*, but slightly lower in hybrids (Supplementary Figure 2C).

For floral morphology, height and width of individual flower display is intermediate in hybrids, with *G. conopsea* producing larger flowers on both axes than those of *G. rhellicani* (Supplementary Figure 3C, D). Similarly, nectar spur length was also found to be intermediate in the hybrids, however closer in length to that of the *G. rhellicani* parent than that of *G. conopsea* (11mm in *G. conopsea*, 4.5mm in hybrids, 1.5mm in *G. rhellicani*) (Figure 1F). For resupination, the angle of labellum rotation which relates to specialised pollination in many orchids (Arditti 2003), the largest angle of rotation was found in *G. conopsea* (average 152.91° ± 17.95°) when compared to *G. rhellicani* (average 8.23° ± 4.37°) which effectively does not resupinate. In accordance with nectar spur length, floral resupination is intermediate but also highly variable in hybrids (average 44.36° ± 20.40°) (Figure 1G). See Tables S3 and S4 for all statistical details of morphology analysis. The morphological differences between these species are expected to result in differential placement of pollinia on pollinators’ bodies, which is expected to limit pollen transfer between them. This also supports the expectation that any interspecific pollen transfer will be in the direction of *G. rhellicani* to *G. conopsea* due to variation in nectar spur length and resupination.

#### Hybrid scent is intermediate, but compound dependent

The *Gymnadenia* orchids in this study were found to produce up to 47 different volatile organic compounds (VOCs), the majority of which being phenylalanine-derived aromatics, also known as benzenoids or phenylpropanoids (26 VOCs) (Figure 2A). These phenylpropanoid VOCs make up a higher proportion of total volatile emission in *G. rhellicani* than *G. conopsea*. In contrast, *G. conopsea* produces higher terpenoid and unknown VOCs as a proportion of total floral volatile emissions. Hybrids do not display global up- or down-regulation of volatile pathways relative to the parent species, but rather relative abundance of volatile patterns in hybrids differ depending on the specific volatile (Figure 2B). For example, whilst absolute phenylethyl acetate production is intermediate in hybrids, phenylethyl alcohol production is more similar to the *G. rhellicani* parent whereas phenylethyl acetaldehyde production is produced in lower levels more similar to *G. conopsea* (Figure 2B). Within the biosynthesis pathway of benzenoid VOCs, these three aromatic VOCs occur within 1-3 biosynthetic steps from the phenylpropanoid precursor, phenylalanine, in turn acting as precursors for each other (Figure 2C). When taken altogether, VOC production is found to be intermediate in hybrids, with significant overlap in patterns of production between species (Figure 2D). See Tables S5, S6, and S7 for all statistical details of floral volatile production analysis.

**Figure 2:**
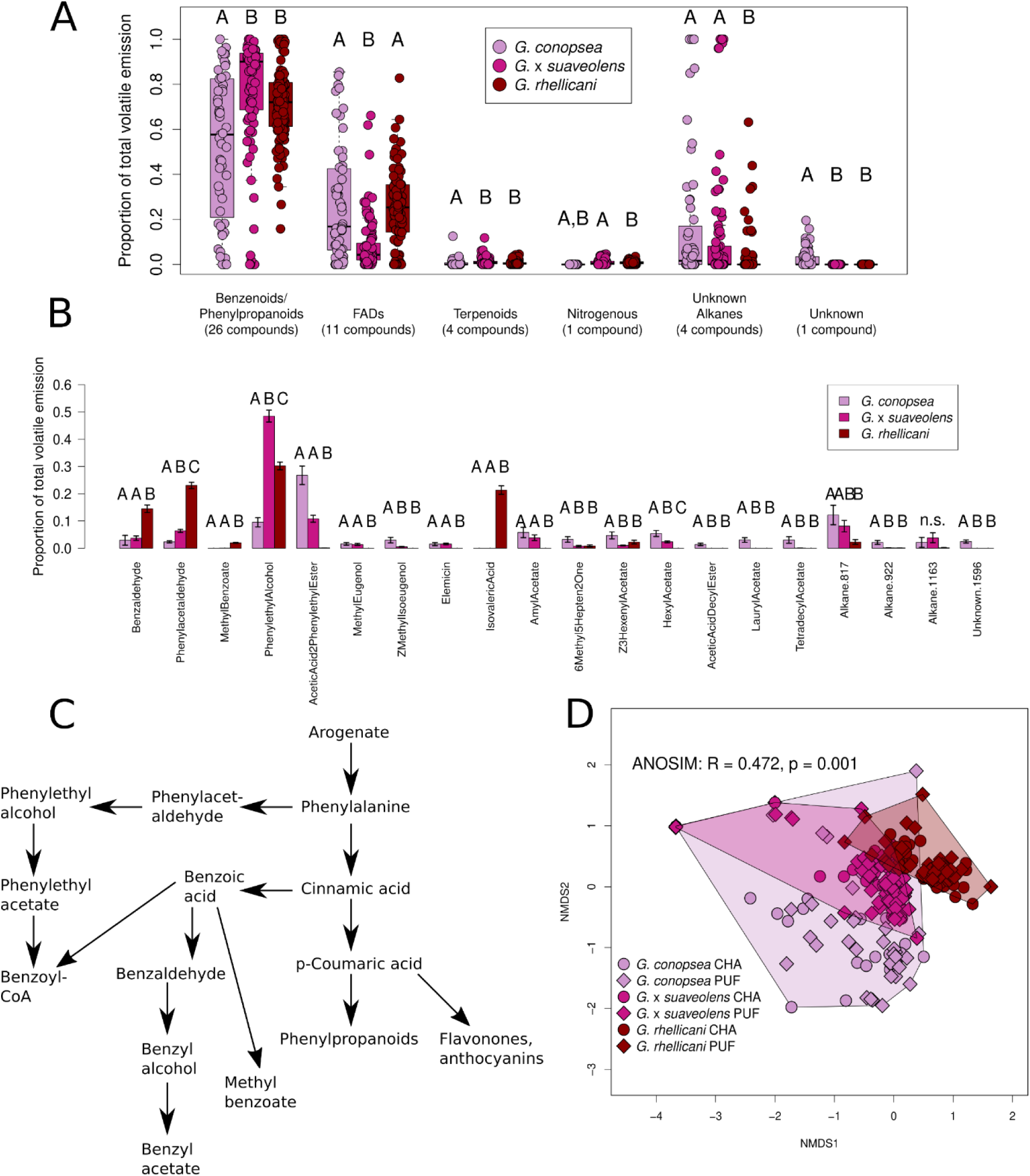
Floral scent of parent and hybrid *Gymnadenia*. (A) Relative (percentage of total) emissions for different volatile compound classes for parent and hybrid orchids. (B) Relative (percentage of total) emissions of major volatiles (those above 1% of total emission in any species) for parent and hybrid orchids. (C) Simplified aromatic volatile biosynthesis pathway (Dudareva et al., 2013) showing the close relationship of phenylacetaldehyde, phenylethyl alcohol, and phenylethyl acetate (upper left) and overall relationship between phenylpropanoid volatiles and anthocyanins as phenylalanine-derived metabolites. (D) NMDS plot of relative (percentage of total) emissions of all floral volatiles for parent and hybrid orchids by site.

#### Hybrid colour has intermediate perception by pollinators

Reflectance spectrophotometry of fresh floral tissue reveals that hybrids are intermediate in floral reflectance between the two proposed parent species, (Figure 3A, B). This is consistent in both the blue and red wavelength peaks, which may be able to distinguish the orchids from the pollinators’ perception (Figure 3C, Supplementary Figure 4A, Table S8) (Telles et al., 2014). When considering hawkmoths (Sphingidae), a Lepidopteran taxon observed to visit *G. conopsea*, none of the *Gymnadenia* species produced peaks in the UV, which would act to repel hawkmoths (Figure 3A) (White et al., 1994). Both hybrids and *G. conopsea* produce a high blue peak that is attractive to hawkmoths such as *Macroglossum stellatarum* (hummingbird hawkmoth) which responds very strongly behaviourally to blue wavelengths (Figure 3C) (Telles et al., 2014). We modelled general Lepidopteran perception as a generic tetrachromatic system (after León-Osper & Narbona 2022), demonstrating that both parents and hybrids fall primarily within the sensitivities of the short- and long-wavelength receptors (Figure 3D), although we note that the long-wavelength sensitive receptor is absent in *Macroglossum* (Telles et al., 2014); the short receptor of *Macroglossum* peaks at 440nm, approximately the wavelength of the “blue peak” (Figure 3C). The intermediate perception of hybrids is also found when considered through the visual perception systems of both bees (using the honeybee *Apis mellifera* as a model) and flies (using the fly *Musca domestica* as a model) (Supplementary Figures 4B, C).

**Figure 3:**
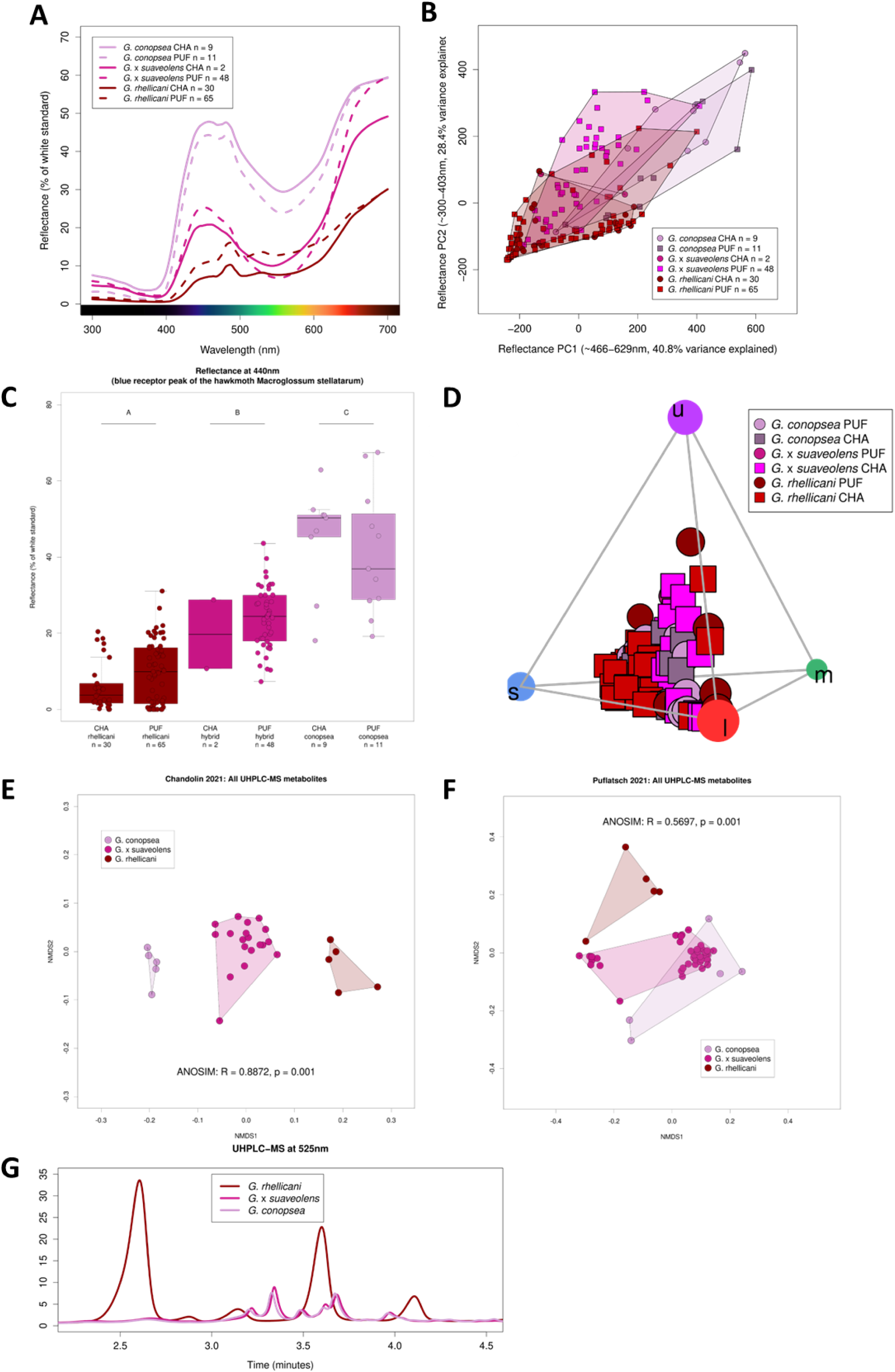
Floral pigment and perception in *Gymnadenia* orchids. (A) Reflectance spectrophotometry curves in the UV-visible spectrum (300-700nm) for each species at each site. Only median values are shown. (B) PCA of the first two principal components of all reflectance spectrophotometry data for each species at each site. (C) Reflectance at 440nm or “human blue” corresponding to the blue photoreceptor peak of the hawkmoth *Macroglossum stellatarum* (Telles et al., 2014). (D) Generic tetrachromat (average avian UV system) visual model plot of reflectance spectrophotometry data for each species at each site; corners represent the four receptors (u = UV; s/m/l = short, medium, and long wavelength receptors respectively). (E, F) NMDS of all floral UHPLC-MS metabolites from Chandolin (CHA) (E) and Puflatsch (PUF) (F) in 2021. (G) UHPLC-MS chromatograms of example parent and hybrid orchids sampled at 525nm.

#### Hybrid pigments are intermediate

Having noted significant visual colour and reflectance differences between the parent species and hybrids, we sought to characterise the anthocyanin production and profiles of these orchids. The prospective parent species, *G. conopsea* and *G. rhellicani*, contain similar levels of total anthocyanins per mg of floral tissue (absorbance at 525nm) to each other, with no significant difference when compared with the hybrids (Supplementary Figure 4D), despite their visually different colours. This is consistent at different sampling locations, with individuals possessing a range of pigment intensities when sampling. It is important to note that all three known pigment morphs of the variable *G. rhellicani* (as described in Kellenberger et al. 2019) were sampled, which may contribute to variability in these values. When looking specifically at all untargeted anthocyanins and anthocyanin-intermediate metabolites, hybrids produce an intermediate metabolite profile when compared to *G. conopsea* and *G. rhellicani* (Figure 3E, F). In contrast to this intermediate total metabolite phenotype, hybrids make a similar final anthocyanin profile as *G. conopsea* as shown in their chromatograms at wavelength 525nm (Figure 3G). This is consistent regardless of sampling year or location. Preliminary individual anthocyanin identification and example raw chromatograms of *G. rhellicani*, *G. conopsea* and 3 hybrids of the same sampling year and location can be found in Supplementary Figure 5.

#### Volatile and pigment modularity differs between parents and hybrids, as well as by population

Considering the relationship between anthocyanins and scent, we sought to uncover whether there are any inter-trait correlations between these two categories of floral phenotypic traits. Interestingly in VOCs, biochemical pathway distance in the aromatics (2014-2016 data only) is correlated with inter-volatile correlation in *G. conopsea* (r = -0.512, p =-10) and hybrids (r = -0.581, p = 10-13), but not in *G. rhellicani* (r = 0.108, p = 0.64). When looking at overall absolute Pearson Correlation Coefficient (PCC) values across the parents and hybrids, we found that these values differed between all three (Supplementary Figure 6A), as did the proportion of significant PCCs (p < 0.05: 9.14% for *G. conopsea*, 10.06% for *G. rhellicani*, and 16.87% for the hybrids, Supplementary Figure 6B, data points below the red line). The hybrids thus seemed to have slightly stronger trait integration than either of the parent species within the volatile and anthocyanin datasets. We also found strong site-specific effects, with absolute PCCs being significantly higher at PUF (mean absolute PCC 0.401) than at CHA (mean absolute PCC 0.311, t = -2.992, df = 1056, p = 0.0028). This suggests that plants at the PUF site overall have stronger integration between volatiles and anthocyanins than plants at the CHA site, perhaps due to differences in local pollinator communities.

### Are the phenotypically intermediate hybrids true F1s?

#### Hybrids and parents do not vary in ploidy levels

We confirmed a lack of interploidy between the proposed parent species with hybrids, concluding that gross-scale ploidy and chromosome count incompatibilities are unlikely to be a reproductive isolating barrier in this system (Figure 4A). We also used this to confirm the viability of using genome proportion analyses for assigning hybrid parentage, as chromosome numbers are shared across the study species.

**Figure 4:**
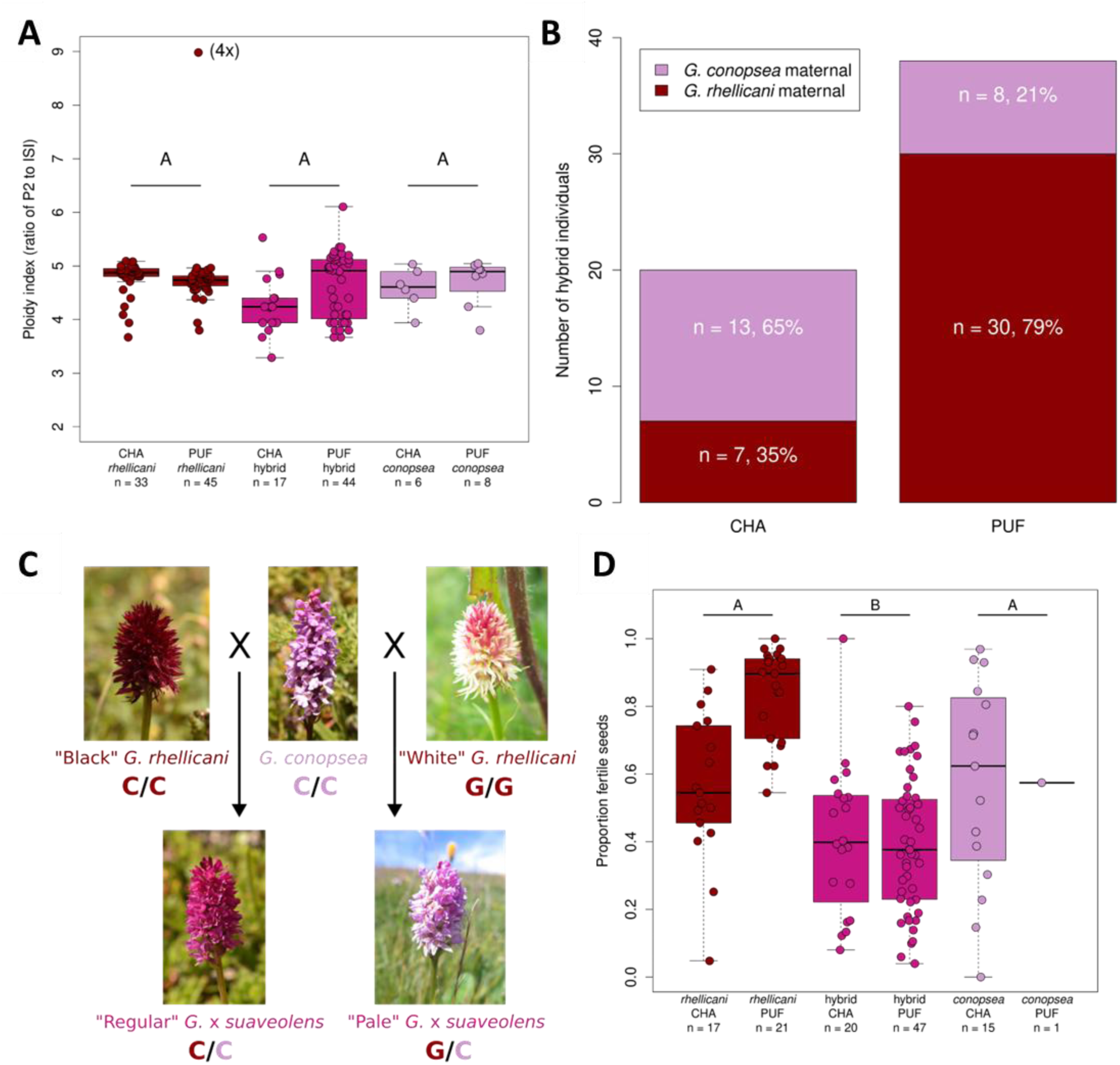
*Gymnadenia* hybrid genetic identity and seed fertility. (A) Ploidy levels of parent and hybrid orchids from the two field sites expressed as the ratio of orchid pollinia tissue peak II (diploid nuclei) to internal standard (*Phaseolus coccinea*) leaf tissue peak I (diploid nuclei). (B)Number of hybrids at each site showing proportion of *G. rhellicani* (red) and *G. conopsea* (pink) maternal parents, p=0.0015 according to Fishers Exact Test. (C) MYB1 Genotypes in the parent and hybrid species of varying colouration (as described in Kellenberger et al., 2019. for *G. rhellicani*). (D) Proportion of fertile seeds per plant for each species at each field site.

#### Hybrids are all F1s and show site-specific bidirectionality in parentage

Using whole genome sequencing, we were able to determine that all hybrids sampled were F1s, resulting from a primary crossing event between the two parent species, as their genomes were all 55/45% split between the parental species. We did not find any evidence of advanced generation hybrids (i.e. F2s) or backcrosses to either parent species. Our work shows bidirectionality in F1 hybrid formation, in which both *G. rhellicani* and *G. conopsea* act as pollen-donor and egg-donor. Interestingly, this bidirectionality is site-specific with the majority of hybrids at CHA being *G. conopsea* maternal and the majority of hybrids at PUF being *G. rhellicani* maternal (Fisher’s Exact Test p=0.0015) (Figure 4B). This demonstrates that either parent species can be pollen or egg parent, but this pattern occurs at different frequencies at the different sites. This also shows that the F1 hybrids cannot be sustained at either site, suggesting a fitness cost when compared to both parents.

Once the parental identities of hybrids were confirmed through sequencing, we used this subset of hybrids to test whether the traits described above were influenced by parentage. For example, we did not find any influence of parentage on hybrid seed set or fertility, with no significant difference in either measurement (total seed set: t = -1.194, df = 31.838, p = 0.241; fertility: t = 0.180, df = 30.493, p = 0.859; total fertile seeds: t = -0.954, df = 34.211, p = 0.347) between *G. conopsea* maternal or *G. rhellicani* maternal hybrids. We also did not find any evidence of parentage influencing VOC production, as individual and global VOC trends are not influenced by the hybrid maternal parent (Supplementary Figure 7).

#### Heterozygous MYB1 and ANS genotypes reflect intermediate hybrid colour

To investigate the genetic basis of the hybrids’ intermediate floral phenotype and F1 heterozygosity, we leveraged our whole genome sequencing and previous knowledge of loci (MYB1 and ANS) controlling anthocyanin production in *G. rhellicani* (Kellenberger et al., 2019; including within-species variation among white, light red and black-red morphs). These three colour morphs correspond to three allelic states at position 663 in MYB1, the most common state being a C, corresponding to black-red floral colour. Here, we sequenced *G. rhellicani* (including all three colour morphs), in addition to co-occurring *G. conopsea* and their uncommon hybrids, neither of which has previously been characterised at these loci. For MYB1, all individuals of *G. conopsea* sequenced at both sampling sites were homozygous for the C allele at position 663. This same SNP is homozygous C/C in most of the hybrids, as expected for F1s between the most common, black-red *G. rhellicani* and *G. conopsea* (Figure 4C, Supplementary Figure 8). When compared to the *G. rhellicani* sequence, we identified an additional SNP conferring a G to A transition at position 688 that is found to be homozygous in *G. conopsea* and heterozygous in the magenta hybrids. This is a non-synonymous SNP, conferring an aspartic acid to asparagine residue shift (Supplementary Figure 8). This further supports the analyses described above confirming that the hybrids are in fact F1s.

Interestingly, the single “pale pink” hybrid we sequenced (which had a notably paler floral colour in the field) is heterozygous (C/G) for the same SNP at position 663, derived from the white *G. rhellicani* sequenced (Figure 4C, Supplementary Figure 8). Other paler hybrids were also observed at Puflatsch in 2015-2016 – these likely also result from a cross between the white *G. rhellicani* morph and *G. conopsea but* were not sequenced.

We also looked at the ANTHOCYANIDIN SYNTHASE (ANS) previously characterised in *G. rhellicani* only by Kellenberger et al., (2019). We found that the *Gymnadenia conopsea* allele has a series of SNPs at the end of the sequence, in addition to a truncated sequence, when compared to the *G. rhellicani* sequence (Supplementary Figure 9A). Once again, we found the hybrids to be heterozygous for the two parental species’ alleles, with a single copy of the full-length sequence and one copy of the truncated sequence inherited from the *G. rhellicani* and *G. conopsea* parent, respectively (Supplementary Figure 9A). This once again confirms the identity of the hybrids as F1s and suggests that the intermediate pigment phenotype of the hybrids may correlate with heterozygosity at these loci, which differ in parental species. Preliminary sequence analysis found the *G. conopsea* allele to indeed confer a truncated protein due to a frameshift mutation (Supplementary Figure 9B), however, we cannot determine at this point whether this truncation confers a change in protein function.

### Do other potential barriers to reproduction explain hybrid rarity?

#### Hybrids make fertile seeds

As detailed above, we did not find evidence that parent species or hybrids varied in ploidy level (Figure 4A), and thus chromosomal incompatibilities arising from differences in chromosome number are unlikely to explain the observed rarity of hybrids and the absence of advanced generations of hybrids.

To understand whether differences in fitness could reflect barriers to reproduction of parents and hybrids, we investigated patterns of seed production and fertility. Given the rarity of hybrids and the lack of subsequent generations of hybrids, we initially hypothesised that hybrids may be sterile, and incapable of producing fertile seeds. However, our results demonstrate that hybrids are capable of producing a substantial proportion of seeds containing embryos (Figure 4D). When considering total volume of seeds, *G. conopsea* was found to produce the greatest number of seeds (average 35,245 ± 33,840 seeds per plant), with *G. rhellicani* (average 3,458 ± 6,689 seeds per plant) and hybrids (average 9,359 ± 15,076 seeds per plant) producing similar amounts (Supplementary Figure 2D). Whilst *G. conopsea* has a higher abundance of total seeds produced, it has a similar proportion of seed fertility (57.08 ± 29.99% fertile seeds) when compared to that of *G. rhellicani* (71.38 ± 22.41% fertile seeds) (Figure 4D). Both parent species were found to have greater seed fertility than that of the hybrids (39.61 ± 20.35% fertile seeds), though hybrids were still capable of producing fertile seeds (Figure 4D). Most hybrid samples produced a substantial proportion of fertile seeds, though some produced few if any fertile seeds. When compared against the abundance of seeds, the fertile hybrid seed set is relatively small (average 3,479 ± 5,725 fertile seeds) in comparison to *G. conopsea* (average 18,216 ± 14,237 fertile seeds) but more similar to that of *G. rhellicani* (average 1,944 ± 3,276 fertile seeds) (Figure 2E). See Table S9 for all statistical details of fitness and fertility. As hybrids can produce seeds and are therefore not considered entirely sterile, the lack of subsequent generations (F2s, etc.) cannot be explained by hybrid sterility alone. Thus, we argue that pollinator-mediated isolation mediated by floral morphology (especially nectar spur length and resupination) and pollen placement of limited shared pollinators is likely a strong isolating barrier in this system.

## Discussion

This study clarifies the context and extent of the formation of rare hybrids between *G. conopsea* and *G. rhellicani* in their zones of overlap in the European Alps, where they share a partially overlapping assemblage of pollinators, thus shedding light on reproductive isolation in this system. We find that hybrids are rare (<1% of the population) and are in fact phenotypically intermediate across nearly all floral traits. Hybrids are largely intermediate between parents in morphology and floral reflectance; however, their anthocyanin profile is the only floral phenotype that replicates that of one parent, in this case the *G. conopsea* parent. Floral scent in hybrids is more variable and not consistent even within closely related volatile organic compounds (VOCs). These largely intermediate hybrid phenotypes reflect exclusively F1 crosses between *G. conopsea* and *G. rhellicani*, as confirmed by whole genome sequencing. No evidence of backcrossing or introgression was found across any study sites, but parentage (which species was pollen or egg donor) varied between sites, contrary to expectations. Additionally, all hybrids were found to be heterozygous at two loci known to impact the distinct anthocyanin pigments observed in parents, reflecting a genetic basis connecting the hybrids’ intermediate phenotypes in floral colour with their genetic inheritance of one allele from each parent species, thus linking phenotype with genotype. Finally, we find little support for several other possible isolating mechanisms, namely that: parents and hybrid offspring do not vary in their respective number of chromosomes and hybrids produce fertile seeds. Thus, we suggest that floral isolation, mediated by distinct floral traits of parents, likely represents an important isolating mechanism between *G. conopsea* and *G. rhellicani* in limiting the formation of uncommon, unfit hybrids.

Here we have characterised the production and perception of the characteristic magenta-pink pigments of *Gymnadenia* hybrids and their aromatic-rich scent profile. Understanding that pigment biosynthesis and scent production are biochemically linked is important when considering that these orchids make both anthocyanin pigments and aromatic volatiles as most of their scent bouquet – both of which are derived from the common precursor amino acid phenylalanine (Dudareva et al., 2013; Li et al., 2024). *Gymnadenia* is a chemically diverse genus with 47 volatiles across both species, the majority of which by total amount are phenylalanine derived. Hybrids globally seem ‘intermediate’ in scent between the two parent species, with no clear pattern of individual volatile inheritance or dominance from either parent species, even for very biosynthetically closely related volatiles such as phenylacetaldehyde, phenylethyl acetate, and phenylethyl alcohol. These volatiles are biochemically all within two reactions of each other (Figure 4C) (Dudareva et al., 2013). Interestingly, many of these phenylalanine-derived aromatics, including phenylacetaldehyde and phenylethylacetate, are known to elicit an electrophysiological response in pollinators of orchids of this group (Huber et al., 2005). Whilst the characterisation by Huber et al., 2005 was initially performed on closely related *Gymnadenia odoratissima* and the previously synonymized *G. densiflora* and *G. conopsea* combined (Huber, pers. comm.), it is important to note that both *G. rhellicani* and *G. conopsea* share major pollinators. Major pollinators of *Gymnadenia* are known to respond to floral scent, including some of the volatiles characterised here, including (but not limited to) benzaldehyde, phenylacetaldehyde, benzyl acetate, phenylethyl acetate and eugenol (Chapurlat et al., 2019; Huber et al., 2005; Jersáková et al., 2010). Further behavioural response analysis through trapping found phenylacetaldehyde to be the most significantly attractive volatile when tested alone (Huber et al., 2005). It is important to note that these previous studies focused on *G. densiflora*, *G. odoratissima*, and *G. conopsea* populations found in Sweden (Chapurlat et al., 2019; Chapurlat et al., 2018) and the Czech Republic (Jersakova et al. 2010), where neither are sympatric with *G. rhellicani*. This study is the first to characterise *G. rhellicani*, *G. conopsea* and their hybrids in their zones of sympatry where floral scent may play a key role in reproductive isolation *via* pollinator attraction in this group.

Interestingly, in this system all species studied produce a high proportion of phenylpropanoid and benzenoid volatiles (Figure 2A). These are derived from phenylalanine and are a result of the shikimate pathway, alongside anthocyanins, suggesting a potential global upregulation of the shikimate pathway in both parental species and hybrids leading to high levels of aromatic volatiles and anthocyanins. The intermediate nature of hybrid scent is mirrored in their total phenylalanine-derived liquid metabolites, reflecting their genetic composition as F1s or primary hybrids. The striking exception to this pattern was the anthocyanin profile of the hybrids at 525nm, which reflect only the *G. conopsea* parent, regardless of whether this parent is maternal or paternal. It remains unclear as to why the hybrids produce the same anthocyanin profile as *G. conopsea* and not *G. rhellicani* or an intermediate anthocyanin profile, as would be expected based on other traits characterised in this study. It is possible that *G. conopsea* alleles are dominant to *G. rhellicani*, either directly in the biosynthesis pathway or higher-level regulators, or that the *G. conopsea* parent confers alleles to the hybrids that further modify the profile through the inclusion of additional enzymatic steps. Alternatively, or in addition, there may be linkage between the anthocyanin pathway genes and genes affecting survival in the hybrids, such as those affecting mycorrhizal recruitment necessary for germination or other survival factors (Li et al., 2021). It would be expected that hybrids may produce biochemically distinct anthocyanins when accounting for their intermediate metabolites. Further genetic analysis will elucidate the role of key metabolism genes and their regulators in the shikimate pathway in both parents and hybrids. This is the pathway which produces both anthocyanins (the most abundant pigment type in this system) and aromatic scent compounds (the most abundant scent components in this system), suggesting it may play an important role in controlling floral signals of these species (Li et al., 2024; Ma & Constabel, 2019; Narbona et al., 2021; Saigo et al., 2020).

We have independently confirmed that at least one of the “pale pink” hybrids is in fact derived from hybridisation between *G. conopsea* and white-morph *G. rhellicani* (Figure 4C). It is worth noting that the white *G. rhellicani* has only been found in Puflatsch and Monte Bondone, Italy, to date (Kellenberger et al., 2019). Moreover, this confirms that this bidirectionality in hybridisation occurs regardless of the colour of the *G. rhellicani* parent, suggesting that pigment-mediated pollinator attraction does not influence the production of these F1 hybrids. Kellenberger et al., (2019) also show that both bees and flies (despite their opposing colour preferences) visited the ‘red’ morph *G. rhellicani* in high frequency, resulting in their increased prevalence and fitness. It is possible that fewer “pale” hybrids are found in Puflatsch simply because fewer white morph *G. rhellicani* are present in the population (ca. 10% of the total *G. rhellicani* surveyed; Kellenberger et al., 2019). Further studies will interrogate the apparent site-specific relationship between anthocyanin biosynthesis genes, in addition to the elucidation of the complete anthocyanin biosynthesis pathway in these orchids, with pollinator perception and observation studies. This will elucidate the role that this biochemical process, alongside the production of aromatic VOCs, has in the attraction of pollinators and thus the evolution and maintenance of these populations.

Consistent with all sequenced hybrids being F1s, we found that the two anthocyanin biosynthesis genes (characterised in *G. rhellicani* by Kellenberger et al., 2019), were heterozygous for the *G. conopsea* and *G. rhellicani* alleles in all sequenced hybrids. However, this does not explain why hybrids only produce *G. conopsea* anthocyanins. Whilst this might suggest that these genes do not play a role in the differences observed in hybrid anthocyanins, it is important to note that at least one of the pale hybrids, only found in Puflatsch to date, has a white-morph *G. rhellicani* parent. This confirms that the white *G. rhellicani MYB1* allele does confer reduced anthocyanin production in hybrids, as they become pale magenta rather than the more typical bright magenta. This implies that the differences in anthocyanin biosynthesis observed between *G. rhellicani* and *G. conopsea* occur during regulation of the biosynthesis pathway, whereby the pale pink hybrid receives less total anthocyanin influx due to the non-functional *MYB1* allele inherited from *G. rhellicani* but ends up with the same profile due to inherited *G. conopsea* elements acting downstream of MYB1. The *ANS* allele in *G. conopsea* does confer an amino acid change leading to a premature stop and truncated protein sequence, when compared to *G. rhellicani*, but it is yet unconfirmed whether this confers a change in protein function. It is likely that both the transcriptional regulator and downstream biosynthesis genes play a significant role in mediating pollinator interactions and acting to prevent the formation of hybrids, as has been observed in *Phlox* (Hopkins & Rausher, 2012).

When considering the hybrid identity via genome proportion analysis and allelic heterozygosity, we were surprised to find all hybrids to be primary hybrids. Whilst making up a small proportion of the population, hybrids do produce seeds containing an embryo, suggesting some level of viability. However, we were unable to test germination or establishment of seedlings (due to challenges of propagation in this study system), which may also act as a barrier to hybrid fitness. Future work should directly evaluate the germination and survivability of hybrid seeds and requirements for germination, such as specific mycorrhizal fungi associations.

Clearly, hybrids are rare. Rather than forming a “hybrid swarm” in this zone of sympatry, the hybrids appear to only emerge in zones where the parent species are found in large abundance, as they have not been seen at otherwise seemingly suitable sites where only one parent species or the other is dominant, with the other species only represented by a few individuals (Byers, pers. comm.). It is possible that F1 hybrids are simply rare enough (<1% of the population at Puflatsch) that we do not observe advanced generation hybrids or backcrosses; alternatively, there may be intrinsic factors rendering these alternatives inviable, or we may have overlooked these plants in the field. However, there have been rare photographic reports of putative backcrosses in the literature (Gerbaud et al., 1999), and their floral colour is different enough from the parent species that they are unlikely to have been overlooked in our comprehensive surveys, where we sampled every hybrid available at each of our field sites.

As demonstrated above, the hybrids produce a small but fertile seed set. It is possible that backcrossed individuals do exist but are no longer phenotypically distinct in the way that *G. x sauveolens* are clearly identifiable in the field and may phenotypically resemble their dominant parent instead. However, backcrossed individuals were not found in the sequencing of parent species, which are genetically distinct, across the two sites. The distinct genetic make-up of the parent species and hybrids found in our sequencing, along with the scarcity of F1 hybrids found at the field sites makes the possibility of significant numbers of phenotypically indistinct backcrossed generations unlikely. It is also possible that factors involving the environment and soil microbiome required for germination of these orchids act as an additional post-zygotic barrier (Li et al., 2021; Lin et al., 2020). In addition to determining that all sequenced hybrids are F1s, we found the formation of hybrids to occur bidirectionally with a significant difference in pollinia transfer direction based on location. This contrasts with the prediction of only unidirectional formation of hybrids - based on expected pollen transfer only from *G. rhellicani* (the short-spurred parent) to *G. conopsea* (the long-spurred parent). This hypothesis, found in the literature, is based on differing parental nectar spur lengths and expected placement of pollinia on pollinator probosces (Claessens & Kleynen, 2011; Vöth, 2000). However, hybrid phenotypes were found to be intermediate and variable regardless of their parentage; as a result, egg or sperm donor identity cannot be identified based solely on the hybrid phenotype.

It is possible that at other sites *G. rhellicani* may hybridise with *G. odoratissima* instead of *G. conopsea*, a hybrid combination known as *G. x heufleri* that is observed occasionally in the zone of sympatry between all three species (Delforge, 2006; Byers, pers. obs.). However, *G. odoratissima* was not found at either Chandolin or Puflatsch in any of the surveyed years (2014-2016, 2021-2023), removing it from consideration as a parent species. We were also able to rule out the sympatric, closely related *Pseudorchis albida* (previously *Gymnadenia albida*) as a parent based on our genomic analyses, although this species was present at both of our study sites (Reinhammar et al., 2002). Further sequencing analysis would investigate any evidence of introgression between the *Gymnadenia* parent orchid species, including *G. odoratissima* and *P. albida*, and whether this potential introgression is reflected by sampling location. Knowing that patterns of directional hybrid formation happen at different frequencies at the two sites would suggest that historical patterns of introgression may also be spatially distinct.

Based on their morphology, in particular the angles of resupination and spur length characterised here, it has been proposed that pollen transfer would only occur in the direction of *G. rhellicani* to *G. conopsea* (Hedrén et al., 2018). This hypothesis of singular directionality would suggest that all hybrids would have a *G. conopsea* maternal parent and *G. rhellicani* paternal parent (Claessens & Kleynen, 2011; Vöth, 2000). However, our work aligns with that of (Hedrén et al., 2018) in which bidirectionality occurs in pollen transfer and both parent species act as paternal and maternal parents, although that study had a much smaller sample size and a single sampling location. Interestingly, our study included two field sites in which the parent species occur in sympatry, 309 kilometres apart. Through whole genome sequencing, we found that this bidirectionality of pollen transfer is site-specific, and statistically significant, with the majority of Puflatsch hybrids being *G. rhellicani* maternal and the majority of Chandolin hybrids being *G. conopsea* maternal. This suggests that pollination of the parent species varies between the two sites. It is possible that different pollinator groups differ in their presence and effective pollination both temporally and spatially, giving rise to hybrids of varying parentage at each sample site (Chapurlat et al., 2015; Meyer et al., 2007; Sletvold et al., 2012). In other words, we could be observing a “geographic mosaic” of effective pollinators leading to a mosaic of maternal identities (Thompson, 2005).

One such pollinator we hypothesise has mediated the formation of hybrids is likely to be moths in the family Zygaenidae (particularly *Zygaena* spp.) as they have been observed on both species. It is also possible that other species have carried pollinia on various points of their bodies and probed the densely packed inflorescences. Pollinia have been observed on the bodies (e.g. under the forelegs or on the ventral metathorax) of two other pollinators, a muscid fly (Muscidae) and a noctuid moth (*Cucullia umbratica*, Noctuidae) collected in 2015 on *G. rhellicani* and *G. conopsea* respectively. This would also provide some explanation as to why the hybrids are so scarce – there may simply be very few opportunities for pollinia to be transferred between the parent species due to their differences in resupination and spur length leading to differential pollinia placement on pollinators. This may also provide an explanation as to why backcrosses or F2 generations have not been observed, as F1s have been shown here to have an intermediate spur length and resupination of flowers. Both nectar spur length and floral resupination can affect placement of pollinia on pollinators, therefore providing a potential mechanism for mechanical isolation between the two parent species. Additionally, the intermediate phenotype of the hybrid in both traits may further inhibit correct placement of pollinia and pollinator identity and thus effective pollination. Both *G. rhellicani* and *G. conopsea* are nectiferous perennials which rely on pollinators of various energetic requirements, for example larger Lepidopterans such as hawkmoths, that have been observed pollinating *G. conopsea* (Vöth, 2000). It is possible this variation in nectar production, alongside nectar spur length, contributes to effective pollinator-mediated reproductive isolation (Jia & Huang, 2022).

Recent work has proposed the concept of pollination systems which are “functional specialist and ecological generalist”, in which a floral structure can be specialised to a pollinator guild but still be pollinated by a range of more generalist pollinators in order to increase fitness (Ohashi et al., 2021; Silló & Claßen-Bockhoff, 2024; Wenzell et al., 2024). It is clear from their morphological and biochemical profiles that these closely related *Gymnadenia* species are functional specialists, in which floral phenotypes are specialised for the attraction of specific functional groups of pollinators (Fenster et al., 2004; Armbruster 2017) . However, historical pollinator observations suggest allowance for more generalist pollinia transfer, particularly in the more ecologically generalist *G. rhellicani*, which would be less common – hence the scarcity of the hybrids. Vital work going forward will also be to study how changing climate and the variation observed in temporality of both flowering time and potential pollinator emergence influences these highly coordinated ecological interactions. *G. rhellicani* is already under threat from rapid environmental change, with all climate change models predicting a reduction of available niches as well as pollinator habitats (Geppert et al., 2020; Kolanowska et al., 2021).

One of the major challenges of this work and exciting avenues for further research is in integrating abiotic factors, in particular climate. Sampling in 2022 was inhibited by extreme weather conditions leading to most plants being senesced or aborting their inflorescences prior to our visit at the typical peak flowering time. Further research will characterise what effect current trends in changing climate have on this system. We are yet to fully understand how changing climate will also influence pollinator emergence. It is possible that with rapidly changing climate- and the possibility for ranges and therefore contact zones of parent species to expand or contract- that hybrids will increase or decrease in frequency, a phenomenon with interesting implications for species interactions. It is apparent from this study that the emergence and maintenance of these hybrids vary geographically, likely as a result of their local pollinator ecology. Studies such as this help to further our understanding of reproductive isolation and speciation at a time where systems such as these charismatic orchids are already under threat from global change.

## Acknowledgements

We thank the relevant authorities (the Dienststelle für Wald, Flussbau und Landschaft, Kanton Wallis, Switzerland and the Amt für Natur, Autonome Provinz Bozen-Südtirol, Italy) for permission to conduct this research and collect samples of protected species at Chandolin and Puflatsch, respectively. The authors would also like to thank Florian Schiestl, Mathias Kneubühler, Richard Lorenz, Barbara Baldan and Sebastiano Nigris and Benjamin Kellenberger for various efforts supporting the conducting of fieldwork from 2014 to 2023. We thank Barbara Keller, Franz Huber, Lena Stransfeld and Karin Gross for technical support, as well as Montana Atwater for sharing additional moth proboscis length reference data. We additionally wish to thank the JIC metabolomics and Bioinformatics platforms for technical support, in particular Paul Brett, Saleha Bakht and Burkhard Steuernagel. We would like to thank Florian Schiestl for support in funding acquisition. We would like to thank the following funding bodies for supporting this work; University of Zurich (UZH) Forschungskredit grant to R.T.K., an EU SP7 PLANT FELLOWS Postdoctoral Fellowship grant (GA-2010-267243) to K.J.R.P.B., and three travel grants from the Claraz Schenkung Zurich (two to R.T.K. and one to K.J.R.P.B.). P.M.S. acknowledges support by the Swiss National Science Foundation (SNF, project 31003A_155943). This work was also supported by a Research Grant from the Royal Society (UK) to Byers (RGS\R1\211279) and start-up funding from the John Innes Centre, including funding from the GEN (BB/P013511/1), BRiC (BB/X01102X/1), and HBio (BB/X01097X/1) Institute Strategic Programmes funded by the UK BBSRC.

## Author Contributions

MN, RTK, PMS, and KJRPB conceived the ideas and designed the methodology. MN, RTK, RC, HG KEW, LH, PMS and KJRPB collected the data. MN, RC, HG, PP, RAK, LH and KJRPB analysed the data. MN, KEW and KJRPB led the writing of the manuscript. All authors contributed critically to the drafts and gave final approval for publication.

## Data and code availability

Sample sizes for all analyses are given in Table S10. The 2014 *G. rhellicani* data from Chandolin and Puflatsch is already published in (Kellenberger et al., 2019). All genome data are available on ENA under accession number PRJEB76404. The R script for seed counting (written by Schlüter & Byers in 2015) is available on GitHub (https://github.com/plantpollinator/CountSeeds) and the shell scripts for sequence analysis are available on GitHub as well (https://github.com/plantpollinator/GymnadeniaGenomics).

