## Supplementary Figures for "Pollinator-relevant floral traits impact bidirectional hybridisation in the orchid genus *Gymnadenia*"

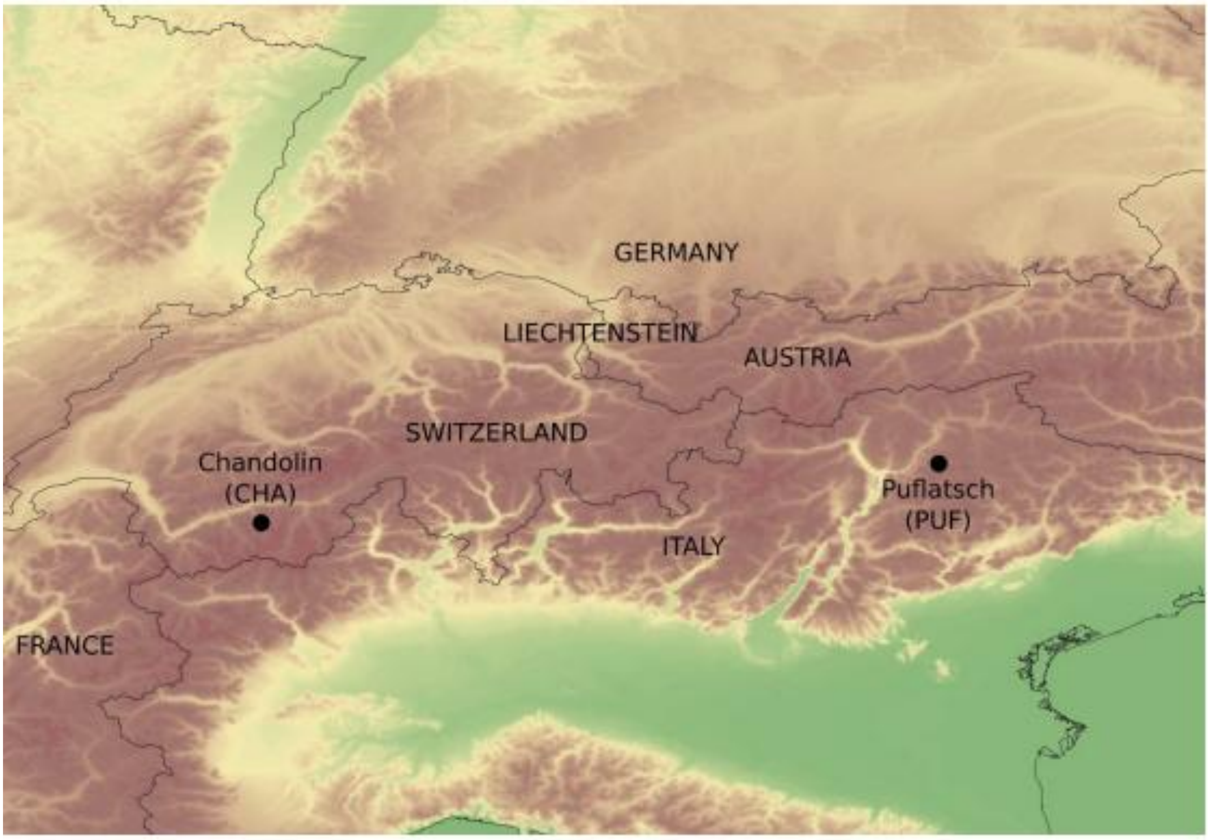


**Supplementary Figure S1: Map showing the two study sites in the elevational context of the Alps.** Chandolin, Switzerland (CHA): 46.255°N, 7.606°E, ca. 2225 masl. Puflatsch, Italy (PUF): 46.555°N, 11.612°E, ca. 2125 masl. The two study sites are separated by approximately 200 miles or 300 km. Raster map elevation data from the GTOPO30 Digital Elevation Model via the European Environment Agency (original data from the United States Geological Survey) with political boundary shape files from GADM, plotted in QGIS

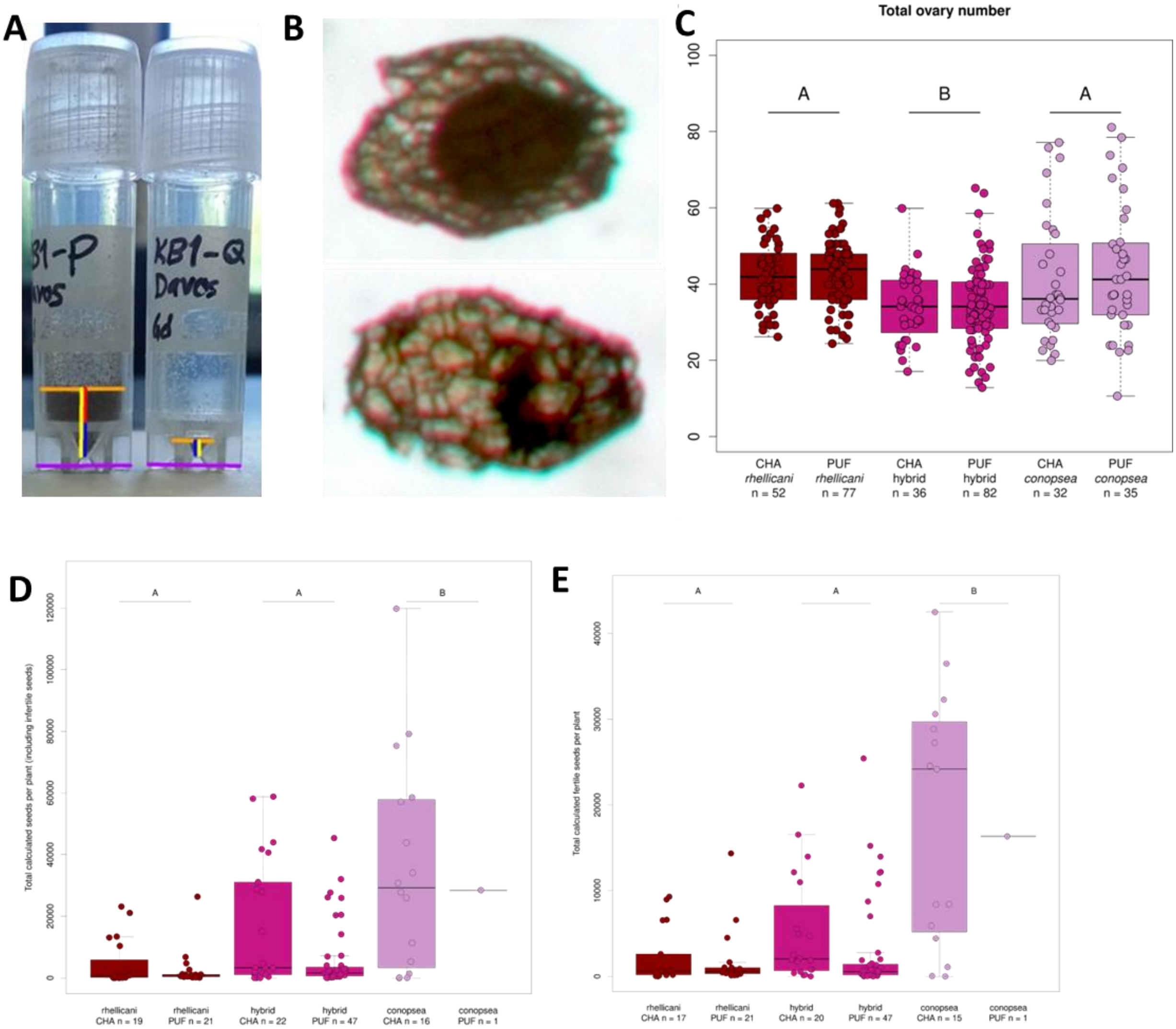


**Supplementary Figure S2: Seed Analysis.** (A)Measurements used to calculate seed volume from photographs taken at a standard distance from the vials, with the mm:pixel ratio calculated from the known dimensions of the vial. Using these measurements, the volume of the vial taken by the seed was calculated, then converted to a total collected seed count using the known volume of *Gymnadenia* seeds (Arditti & Ghani, 2000) and then scaled up to the total plant seed count using the ratio of collected fruits to total ovaries. (B) Example of seed fertility scoring criteria – the upper seed is fertile, with a well-formed rounded embryo. The lower seed is infertile. (C) Total number of potential ovaries (total flowers per inflorescence). (D)Total number of seeds per species per site. (E) The vector product of total seeds (Supplementary Figure 2C) and fertility (Figure 4C) to get the total number of fertile seeds.

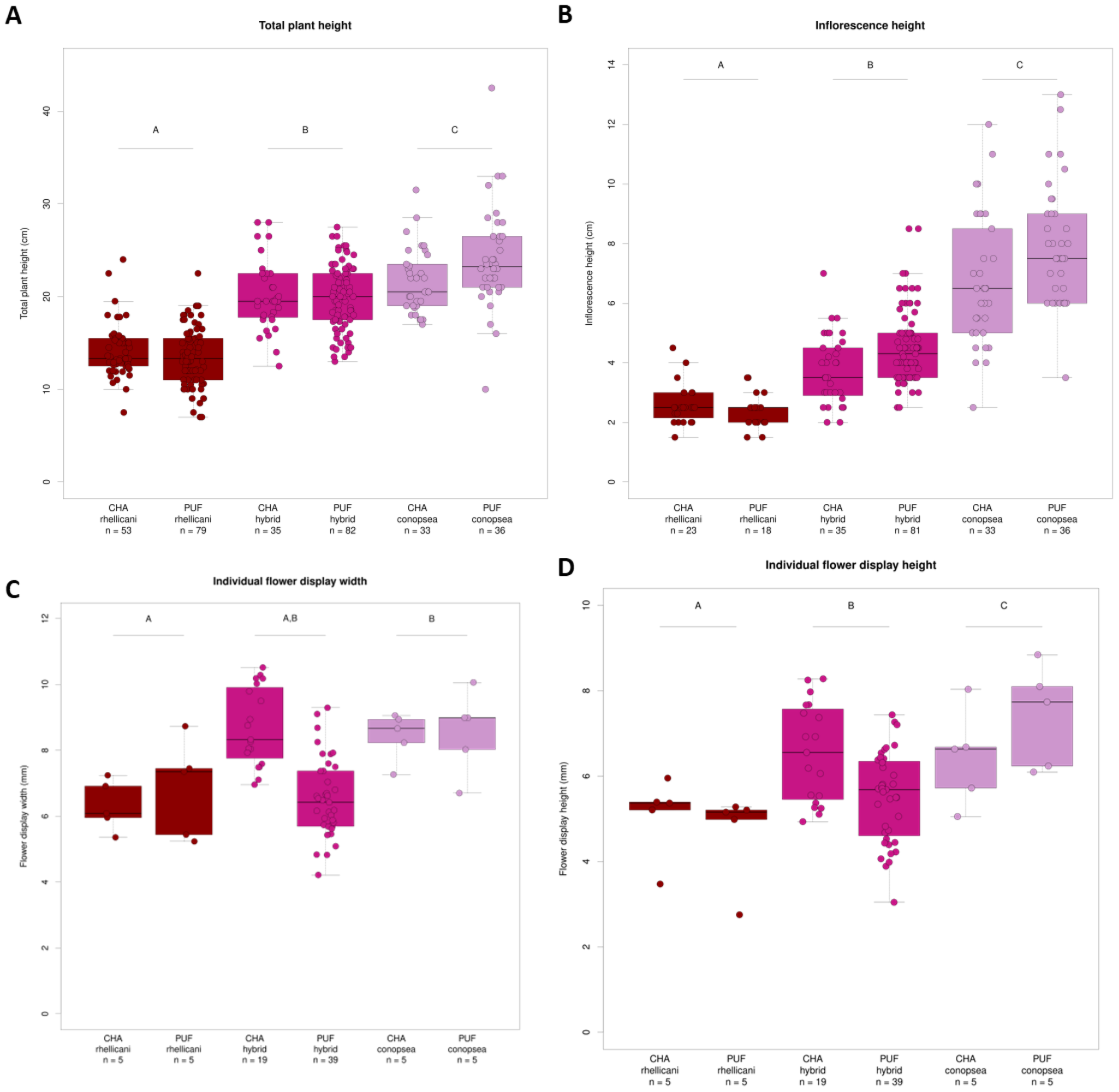


**Supplementary Figure S3: Morphology.** (A) Total plant height in centimetres by species and site. (B) Total inflorescence height in centimetres by species and site. (C) Individual flower display width (the distance between the tips of the lateral tepals) in millimetres by species and site. (D) Individual flower display height (the distance between the tip of the labellum and the opposite tepal) in millimetres by species and site.

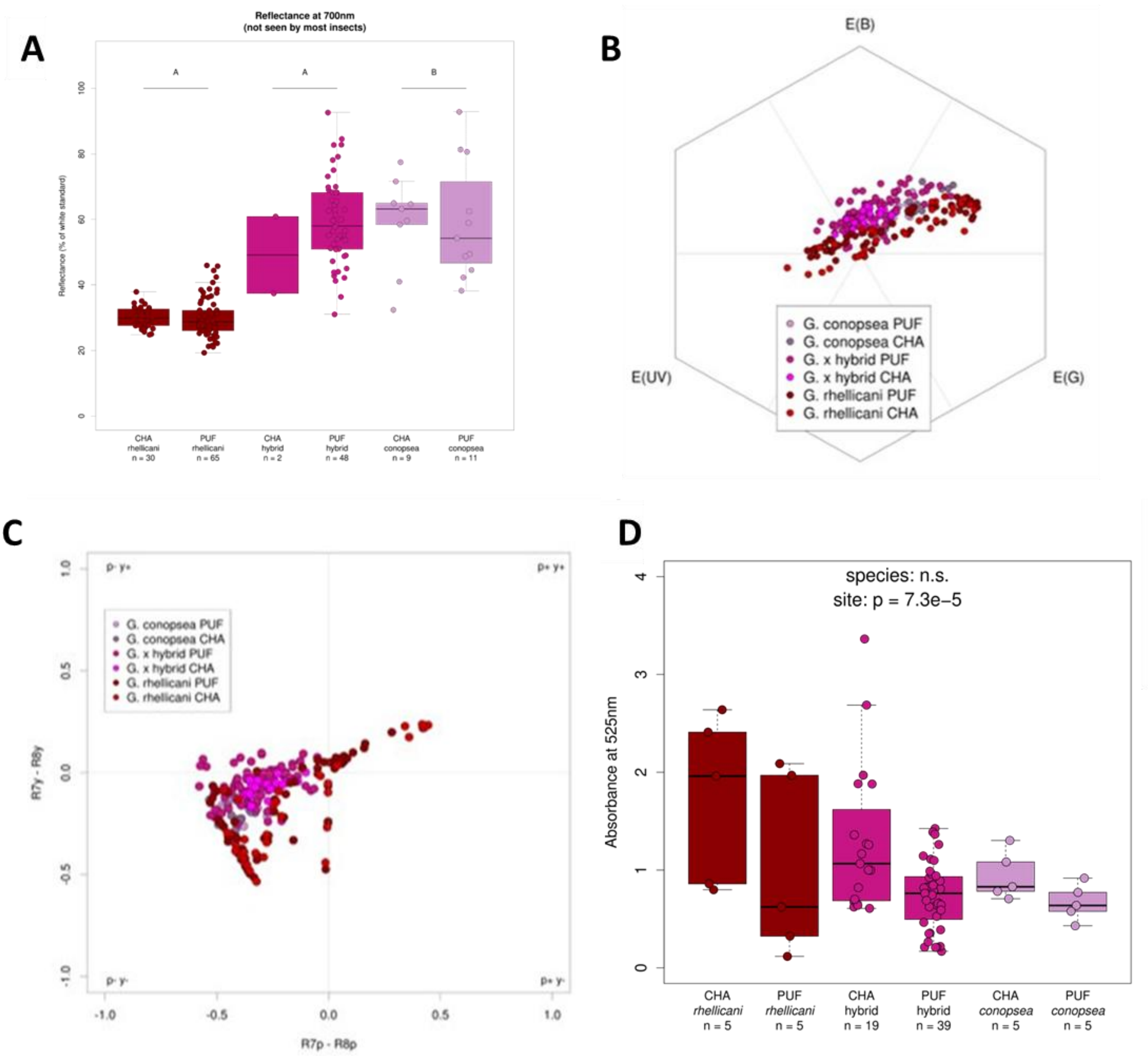


**Supplementary Figure S4: Perception & Pigment.** (A)Reflectance at 700nm (human red, not visible to most insects). (B) *Apis mellifera* (honeybee) hexagon visual model plot of reflectance spectrophotometry data for each species at each site; the centre point represents the achromatic centre. (C) *Musca domestica* (house fly) visual model plot of reflectance spectrophotometry data for each species at each site; only points in different quadrats are distinguishable. (D) Total floral anthocyanins measured via absorbance at 525nm.

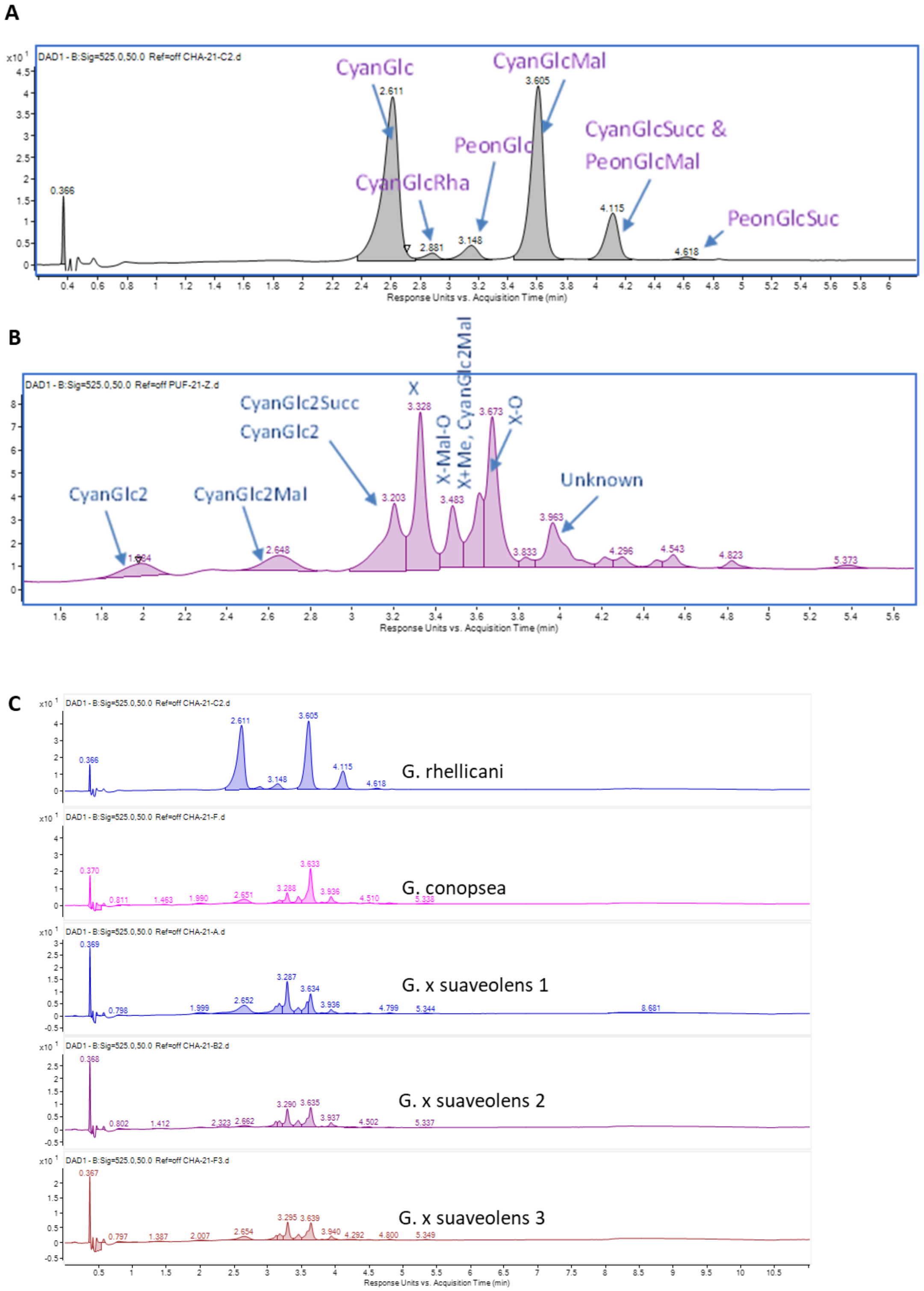


**Supplementary Figure S5: Anthocyanin Identification.** (A)Preliminary anthocyanin identification of *G. rhellicani* from UHPLC-MS data. (B) Preliminary anthocyanin identification of *G. conopsea* from UHPLC-MS data. Cyan = cyanidin, Peon = peonidin, "X" is a complex Orchicyanin-like anthocyanin, Glc = glucoside, Succ = succinyl, Mal = malonate, O= oxalate. (C) Raw chromatograms of *G. rhellicani*, *G. conopsea* and three separate individuals of *G. x suaveolens* showing peaks identified in (A) & (B).

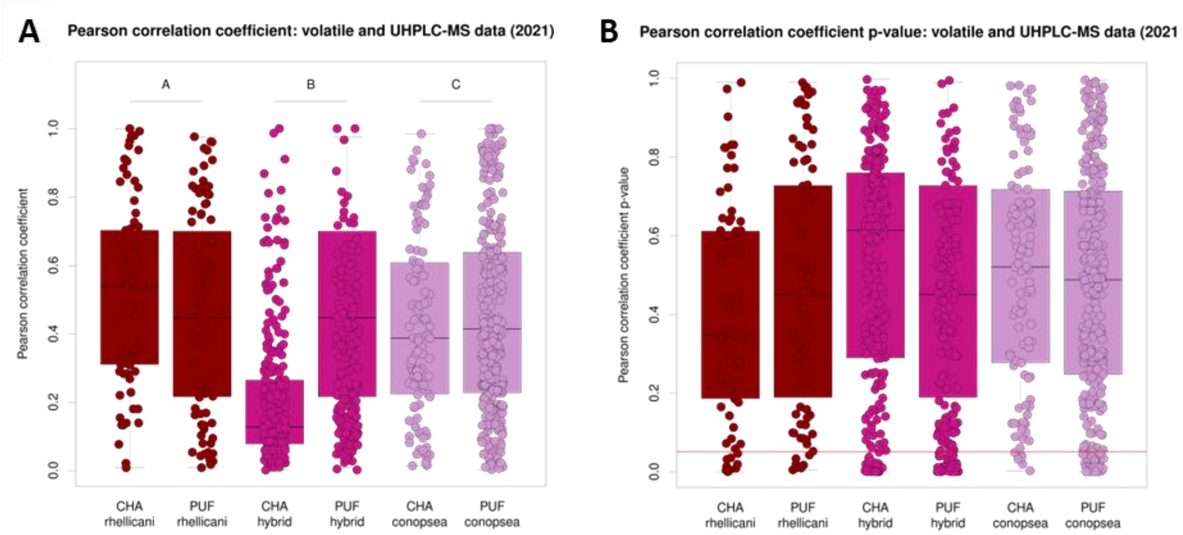


**Supplementary Figure S6: Anthocyanin & Volatile Correlations.** Volatile and UHPLC-MS metabolite correlations from 2021 data. (A) Boxplots of the absolute values of Pearson Correlation Coefficients by species and site. (B) Boxplots of the p-values of the Pearson Correlation Coefficients in (A), with a red line at the p = 0.05 threshold.

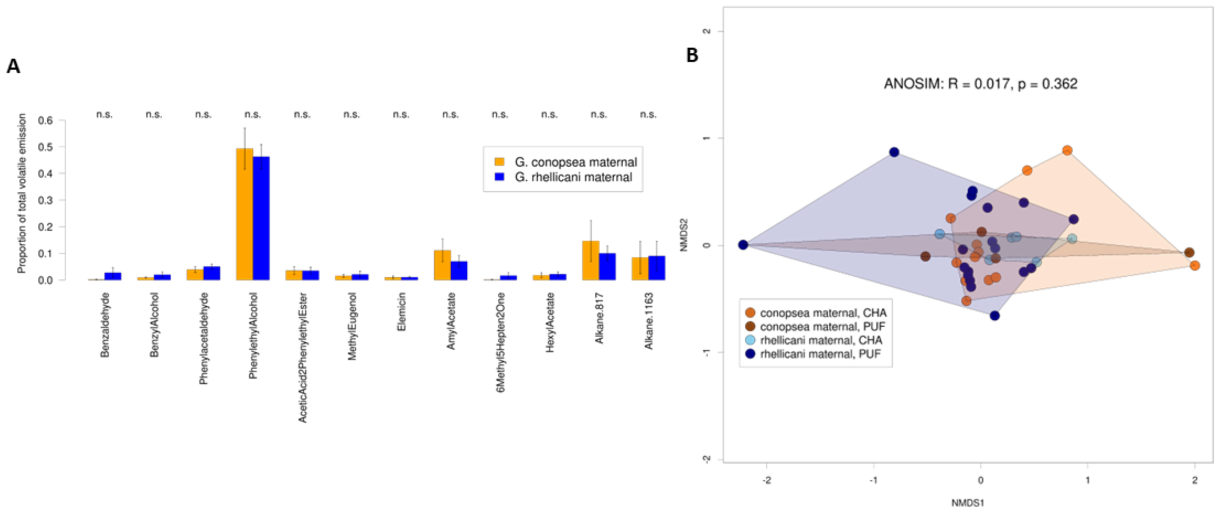


**Supplementary Figure S7: Scent by Parentage.** (A) Major hybrid floral volatile compounds depicted for both maternal identities. (B) NMDS plot of all floral volatile compounds in hybrid orchids separated by maternal identity and site.

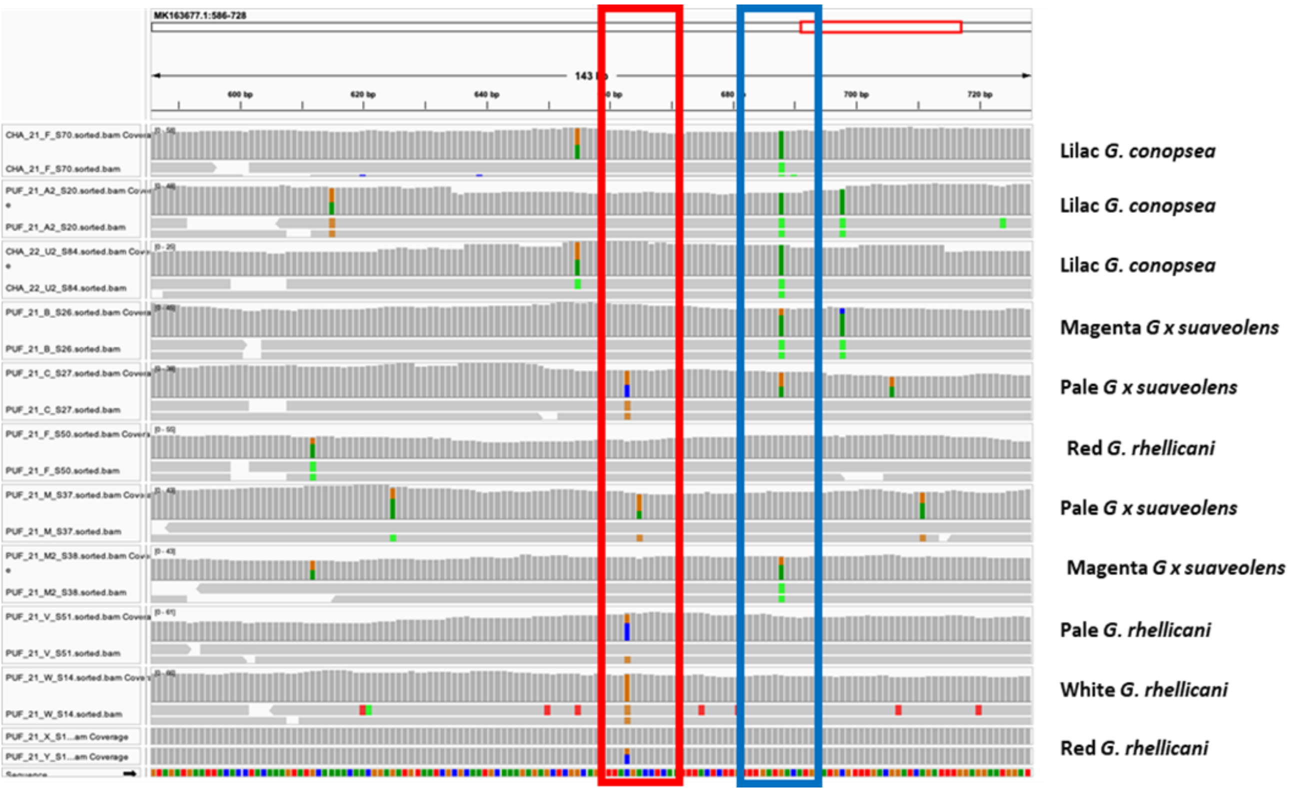


**Supplementary Figure S8: MYB1 Genotypes.** IGV screenshot of *MYB1* sequences. SNP 663, the diagnostic SNP from (Kellenberger et al., 2019) is highlighted in the red box. The blue box indicates the SNP highlighting hybrid heterozygosity at position 688. Species, floral colour, and genotype are indicated on the right. *Gymnadenia rhellicani* colour categories from (Kellenberger et al., 2019).

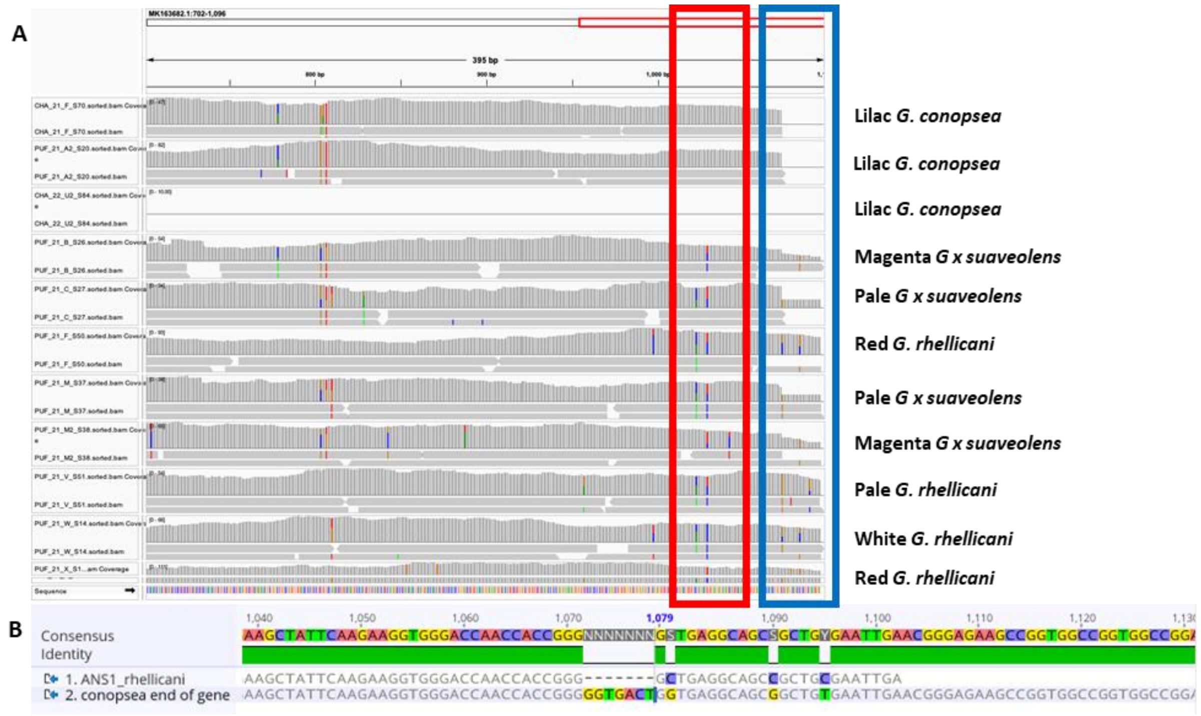


**Supplementary Figure S9: ANS Genotypes.** (A) IGV screenshot of *ANS* (*Anthocyanidin Synthase*) sequences. ANS is truncated (blue box) and has a SNP leading to a premature stop codon (red box) in *G. conopsea*. (B) AlphaFold screenshot demonstrating premature truncation of ANS in *G. conopsea*.
